# A single cell and spatial genomics atlas of human skin fibroblasts in health and disease

**DOI:** 10.1101/2024.12.23.629194

**Authors:** Lloyd Steele, Chloe Admane, Keerthi Priya Chakala, April Foster, Nusayhah Hudaa Gopee, Simon Koplev, Pasha Mazin, Bayanne Olabi, Kenny Roberts, Catherine Tudor, Elena Winheim, Karl Annusver, Donovan Correa-Gallegos, Agnes Forsthuber, Luc Francis, Sophie Frech, Clarisse Ganier, Thomas Layton, Yingzi Liu, Hao Yuan, Johann E. Gudjonsson, Beate M. Lichtenberger, Satveer Mahil, Jagdeep Nanchahal, Edel A O’Toole, Maksim Plikus, Yuval Rinkevich, Emanuel Rognoni, Catherine Smith, Sarah A Teichmann, Maria Kasper, Mohammad Lotfollahi, Muzlifah Haniffa

## Abstract

Fibroblasts are critical cells that shape the architecture and cellular ecosystems in multiple tissues. Understanding fibroblast heterogeneity and their spatial context in health and disease has enormous clinical relevance. In this study, we constructed a spatially-resolved atlas of human skin fibroblasts from healthy skin and 23 skin disorders. We define 6 major skin fibroblast populations in health and a further three skin disease-specific fibroblast subtypes, and demonstrate the fibroblast composition in different types of skin disease. We characterise a human-specific fibroblastic reticular cell (FRC)-like subtype in the skin perivascular niche and postulate their origin from prenatal skin lymphoid tissue organiser (LTo)-like cells. We also show that inflammatory myofibroblasts (*IL11*+*MMP1*+*CXCL5*+*IL7R+*) are a conserved fibroblast subtype in inflammatory disorders and cancers across multiple human tissues. We provide a harmonised nomenclature for skin fibroblasts that integrates previous findings from human skin and other tissues.

## Main text

Fibroblasts are crucial cells for shaping tissue architecture and immune cell niches^1,2^. Heterogeneity within fibroblast subpopulations has been difficult to study due to a lack of unique surface markers and their shift to activated phenotypes during *in vitro* culture^1,2^. Single-cell RNA sequencing (scRNA-seq) and spatial genomics technologies have overcome these challenges^3^, enabling the dissection of fibroblast heterogeneity in human tissues.

Fibroblast-mediated processes, such as scarring and maintenance of immune cell niches, are observed across multiple tissues, suggesting that fibroblasts deploy common mechanisms across organs. Pioneering cross-tissue fibroblast atlases have begun to elucidate common fibroblast states across different human organs^4–7^. However, these studies have focused on few diseases (1-3 diseases per organ), without comprehensive definition of fibroblast subtypes in healthy states or their spatial localisation within tissues. Consequently, the fibroblast composition of a human organ and how this changes across a range of disease contexts, including immune-mediated diseases, cancer, and fibrosis/scarring, is still unclear.

In this study, we used published large-scale scRNA-seq datasets of normal human skin and 23 diseases, newly generated spatial transcriptomics (whole transcriptome Visium and imaging-based single cell resolution Xenium), and machine learning models to construct a high-resolution atlas of more than 350,000 adult human skin fibroblasts. We provide a consensus annotation of skin fibroblasts based on gene expression profile and spatial location within human skin, contextualised using findings from across human tissues. We characterise the fibroblast compositional signatures in specific skin disease phenotypes and explore the origin of disease-specific fibroblast subtypes. We also reveal unique prenatal skin lymphoid tissue organiser (LTo)-like cells that likely give rise to the adult skin fibroblastic reticular cells (FRC)-like cells in the perivascular niche. Our study harmonises human skin fibroblast nomenclature with the aid of a centralised community annotation platform (CAP) (https://celltype.info/project/388) for the Human Skin Cell Atlas^8^. Our scRNA-seq and spatial datasets resources are freely available for download and interactive data exploration through the following integrated link (to be added upon publication)

## Results

### Human skin fibroblast subtypes are located in distinct tissue microenvironments

We integrated 2.1 million cells from scRNA-seq data of adult human skin, comprising 32 datasets and 251 samples (Supplementary Table 1)^9–40^ using scVI^41^. Fibroblasts were selected based on expression of canonical marker genes, including *PDGFRA*, *DCN*, and *LUM* (Extended Data Fig. 1a). In total, 357 276 high-quality fibroblasts were included post-quality control (Fig. 1a and Methods).

**Figure 1.**
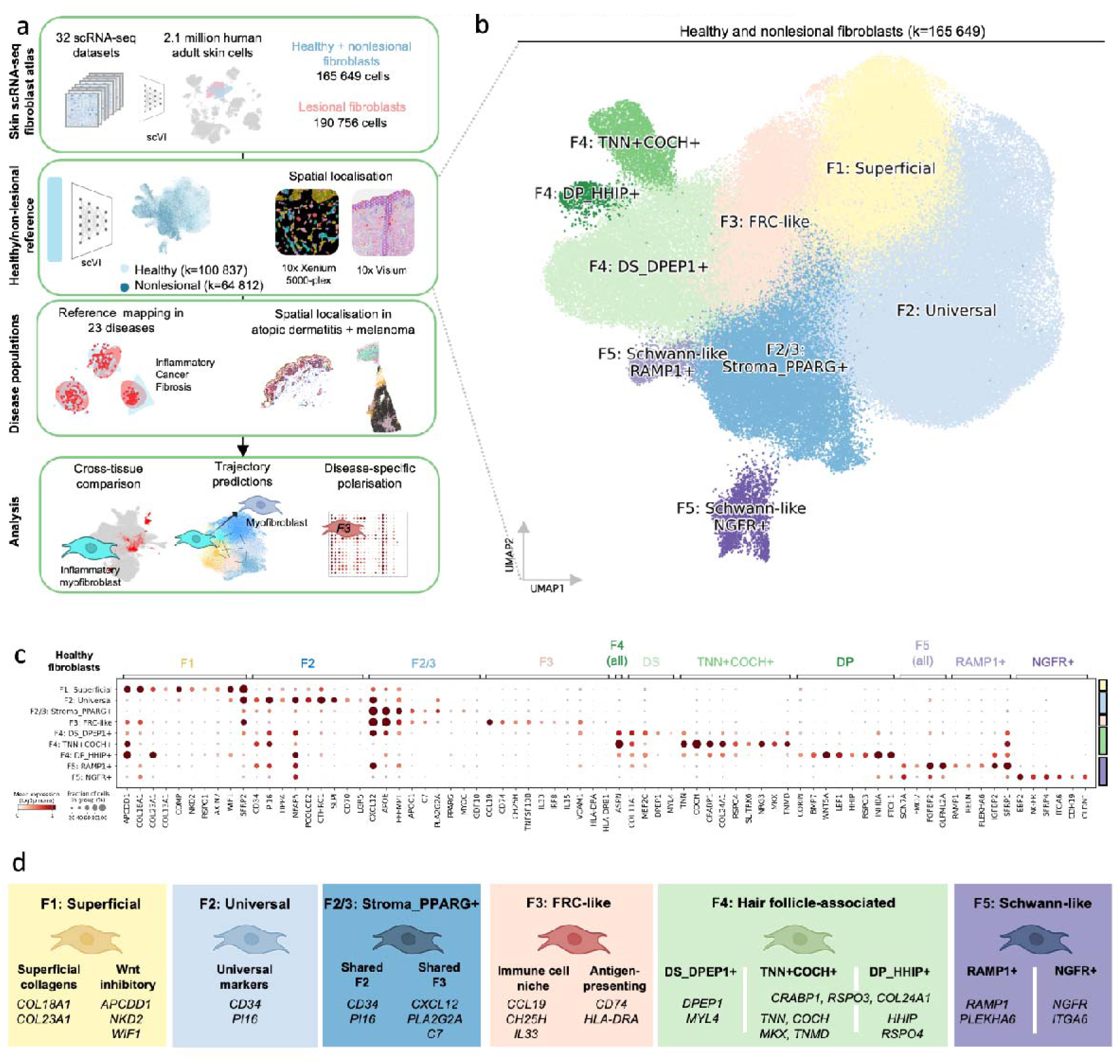
Identification of fibroblast subtypes in healthy skin. a) Overview of study methodology, including skin atlas integration to delineate fibroblasts, construction of a healthy/non-lesional reference, mapping of 23 diseases to the healthy reference atlas, and downstream analysis for cross-tissue comparison. b) UMAP of healthy and non-lesional skin fibroblasts coloured by fibroblast subtype. DS: Dermal sheath fibroblasts. DP: Dermal papilla fibroblasts. c) Dotplot of marker gene expression for healthy fibroblasts. “All” indicates a marker for a general population, but which contains subtypes. See Extended Data Fig 2a for additional differentially expressed genes for fibroblast subtypes. d) Summary of skin fibroblast subtypes in healthy steady-state tissue.

In healthy and histologically normal (non-lesional) skin, we identified six major fibroblast clusters (Fig. 1b,c and Extended Data Fig. 2a). These fibroblast subtypes were observed across multiple datasets and patient samples (Extended Data Fig. 1b-f). To define the spatial location of fibroblast subtypes, we performed array-based spatial transcriptomics (10x Visium) on healthy skin and inflamed atopic dermatitis skin, with deconvolution of spots to individual cell types using cell2location (Fig. 2a-b and Extended Data Fig. 3)^42^. As an orthogonal single-cell resolution spatial method, we used imaging-based spatial transcriptomics (10x Xenium) with a 5000-plex gene panel (Fig. 2c-d and Extended Data Fig. 4). These complementary spatial genomics methods validated the presence of each of the six fibroblast subtypes and revealed their distinct microanatomical locations within human skin.

**Figure 2.**
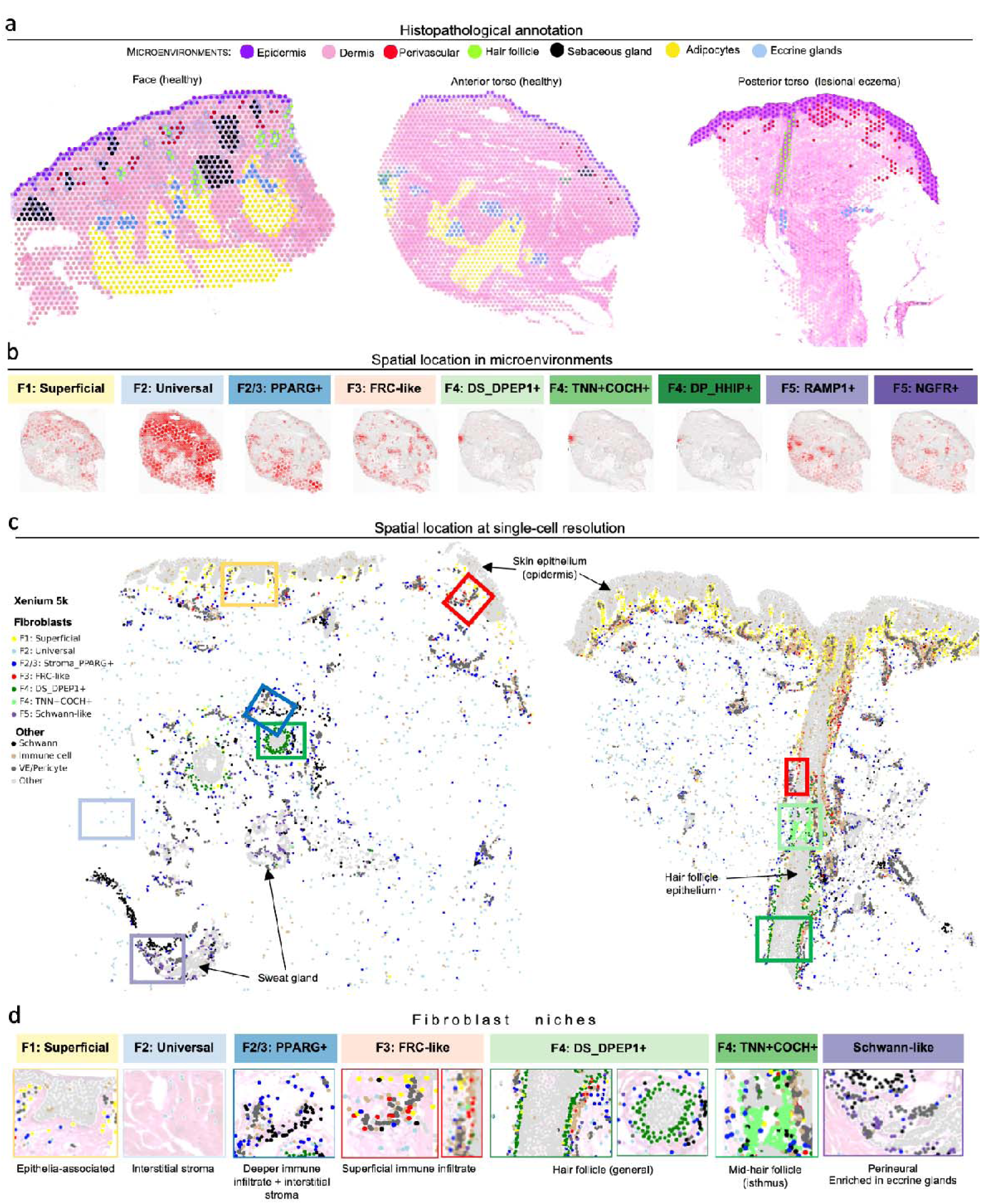
Skin fibroblasts occupy unique spatial and functional niches. a) Histopathological annotation of tissue microenvironments. Each spot is 55 micrometres. b) Spatial location of fibroblast subtypes in microenvironments, predicted by cell2location abundance predictions (10x Genomics Visium), in a single section of healthy human skin. See Extended Data Fig 3a for predictions for all 3 tissue sections. c) Spatial location of fibroblasts at single-cell resolution (10x Genomics Xenium 5000-gene panel) for skin sections from non-lesional skin of atopic dermatitis, coloured by cell type. d) Summary of fibroblast niches: Xenium cell types overlying H&E stained image (manual approximations. See Extended Data Fig. 4e for original H&E images). H&E: Haematoxylin and eosin.

Two of the six fibroblast populations were uniformly present throughout the dermis in the skin sections but were localised to different skin depths. The first population, *F1: Superficial*, showed a clear localisation adjacent to the skin epithelium within the papillary dermis (Fig. 2c,d). Skin fibroblasts in this region have previously been called *Papillary* fibroblasts^33,43–45^. *F1: Superficial* expressed genes encoding superficial dermal collagens (*COL13A1*, *COL18A1*, *COL23A1*) and Wnt signalling inhibitors (*APCDD1*, *WIF1*, and *NKD2*) (Fig. 1c).

A Wnt-mediated, synergistic interplay between superficial dermal fibroblasts and basal epithelial cells has been reported to maintain cellular identity^44^, suggesting a key role for *F1: Superficial* fibroblasts in epidermal homeostasis.

The second population, *F2: Universal,* was located deeper in the skin, interspersed between collagen fibres in the reticular dermis (Fig. 2c,d and Extended Data Fig. 4f). *F2: Universal* are equivalent to previously described skin *Reticular* fibroblasts (Extended Data Fig. 2b,c)^33,46,47^. This population was characterised by high expression of marker genes of *Universal PI16+* fibroblasts (*PI16*, *CD34*, and *DPP4)*^4^. *Universal* fibroblasts are found in many human tissues and are postulated to represent a precursor fibroblast cell state^48,495,6^. Fascial fibroblasts are also reported as potential progenitor cells in mice^50^, but are found deep beneath the fat layer of the skin and are often not captured from skin biopsies. Nevertheless, to understand their relationship to *F2: Universal*, we included fascial fibroblasts obtained from human non-lesional palmar fascia (specific hand anatomical site with minimal fat) in an additional integration (see Methods)^16^. Fascial fibroblasts formed a subset of *F2: Universal* fibroblasts (*F_Fascia*), expressing *F2: Universal* marker genes (*PI16*, *CD34*, and *DPP4)*, in addition to known fascial markers from mice, such as *FGF18* and *CCN3* (Extended Data Fig. 1g,h)^51,52^. These results suggest that *F2: Universal* in skin and fascial fibroblasts are transcriptomically similar, and may have a progenitor role.

The remaining fibroblast subsets were more focal in localisation, being associated with vascular or adnexal structures. We therefore leveraged haematoxylin and eosin (H&E) staining of skin to illustrate these microenvironments and define the fibroblast subtypes associated with them (Fig. 2a and Extended Data Fig. 3b-f). The *F3: FRC-like* population expressed genes associated with specialised fibroblasts, termed fibroblastic reticular cells (FRCs), found in lymphoid organs/structures (Extended Data Fig. 1i,j)^53^. *F3: FRC-like* fibroblasts were enriched in superficial perivascular regions in skin (Extended Data Fig. 3a-d), and at a single-cell level localised with immune cells both in the superficial perivascular region and adjacent to the hair follicle (Fig. 2c,d), where immune cells are known to reside in healthy skin^54^. The genes highly expressed by *F3: FRC-like* fibroblasts are known to attract and compartmentalise immune cells (*CCL19*, *CXCL12, CH25H*), maintain immune cell survival and function (*IL33*, *IL15*, *TNFSF13B, VCAM1*), and enable antigen presentation (*CD74,* MHC-II molecules)^53^.

*F2/3: Stroma_PPARG+* fibroblasts were enriched in deeper perivascular immune infiltrate regions and the interstitial stroma (Fig. 2b-d and Extended Data Fig. 4g). This population shared a subset of marker gene expression with both *F2: Universal* (*CD34*, *PI16*) and *F3: FRC-like* (*CXCL12*, *APOE, C7*, *PLA2G2A*) fibroblasts, but did not express other key *F2/F3* markers (*CCL19*, *SLPI*, *PCOLCE2*). Other studies have also reported a *CXCL12*^hi^*CCL19*^lo^ fibroblast population in human skin^49,55^. *F2/3: Stroma_PPARG+* showed elevated expression of *PPARG* and *SFRP1* (Fig. 1c and Extended Data Fig. 4a). The capability to differentiate into adipocytes is characteristic of reticular fibroblasts (equivalent to *F2: Universal*)^44,46,56^, and the expression of *PPARG* and *APOE* suggests that this population is a subset of the *F2: Universal*/reticular fibroblast lineage that is enriched in pre-adipocytes.

*F4: Hair follicle-associated* fibroblasts encompass 3 subpopulations that are transcriptionally distinct and associated with specific regions of the hair follicle. The first is the dermal sheath (DS) fibroblast population (*F4: DS_DPEP1+*), which has been well-characterised previously (*DPEP1*, *MEF2C*, and *MYL4*)^9^. We now identify a *F4: TNN+COCH+* subset localised to the isthmus (mid-hair shaft) (Fig. 2c and Extended Data Fig. 4c), expressing *CRABP1/COCH/RSPO4* and tendon-associated genes (*MKX*, *TNMD*)^57^. Finally, we identify a fibroblast population uniquely expressing *CRABP1/COCH/RSPO4* and the dermal papilla marker genes *CORIN*, *HHIP*, *RSPO3,* and *LEF1*^58–60^, which we refer to as *F4: DP_HHIP+*. However, as the dermal papilla was not specifically captured in the tissue section analysed by H&E and Xenium (Extended Data Fig. 4d-e), we were unable to validate these cells in situ.

*F5: Schwann-like* fibroblasts (*F5: RAMP1+* and *F5: NGFR+*) were enriched near eccrine glands and related to nerve innervation (Fig. 2b-d and Extended Data Fig. 3f and 4h). *F5: Schwann-like* fibroblasts co-expressed Schwann cell-markers, including *SCN7A* and *ITGA6*. Fibroblasts are known to be located in the endoneurium and perineurium of nerve fibres^61^, and similar fibroblasts have been reported in scRNA-seq studies both in human lung (termed “nerve-associated fibroblasts”)^62^ and skin tissue (termed “Schwann-like fibroblasts”)^63^.

To distinguish *F5: Schwann-like* fibroblasts from Schwann cells and pericytes, we included these populations in our integrated dataset. We observed that Schwann cells and *F5* fibroblasts formed separate clusters and were distinguished by the differential expression of fibroblast markers in *F5* (*PDGFRA*, *LUM*) and *SOX10* and *S100B* in Schwann cells (Extended Data Fig. 1h,k-l). Pericytes were distinguished by *RGS5* expression (Extended Data Fig. 1l).

Overall, we provide a new framework for healthy human skin fibroblast annotation based on gene expression profiles (Fig. 1c,d) and spatial location (Fig. 2d) that integrates previous fibroblast descriptions in the skin and across human tissues^4,55,62–64^ (Extended Data Fig. 2d)^33,44,47,49,65^. Our findings of transcriptionally defined fibroblast subtypes in distinct skin microanatomical locations suggest a role for regional fibroblasts in supporting distinct niche functions within the skin ecosystem.

### Skin disease-adapted and -specific fibroblasts

We next sought to identify how fibroblast composition differs in disease states in skin. We used scPoli; a deep learning model for scalable integration and identification of novel cell states in single-cell transcriptome data^66^. This approach involves constructing an integrated healthy/non-lesional skin atlas (reference) comprising multiple datasets to subsequently map disease (query) single-cell datasets using transfer learning (Fig. 3a)^67^. This permits the contextualization of the disease (query) data to the healthy (reference) data and automated cell annotation. Importantly, the model identifies disease cells that do not map to the healthy reference, thereby highlighting disease-related fibroblast populations. Using this approach, we mapped 190,756 fibroblasts from skin diseases to our healthy/non-lesional *F1-F5* fibroblast (including *F_Fascia*) reference.

**Figure 3.**
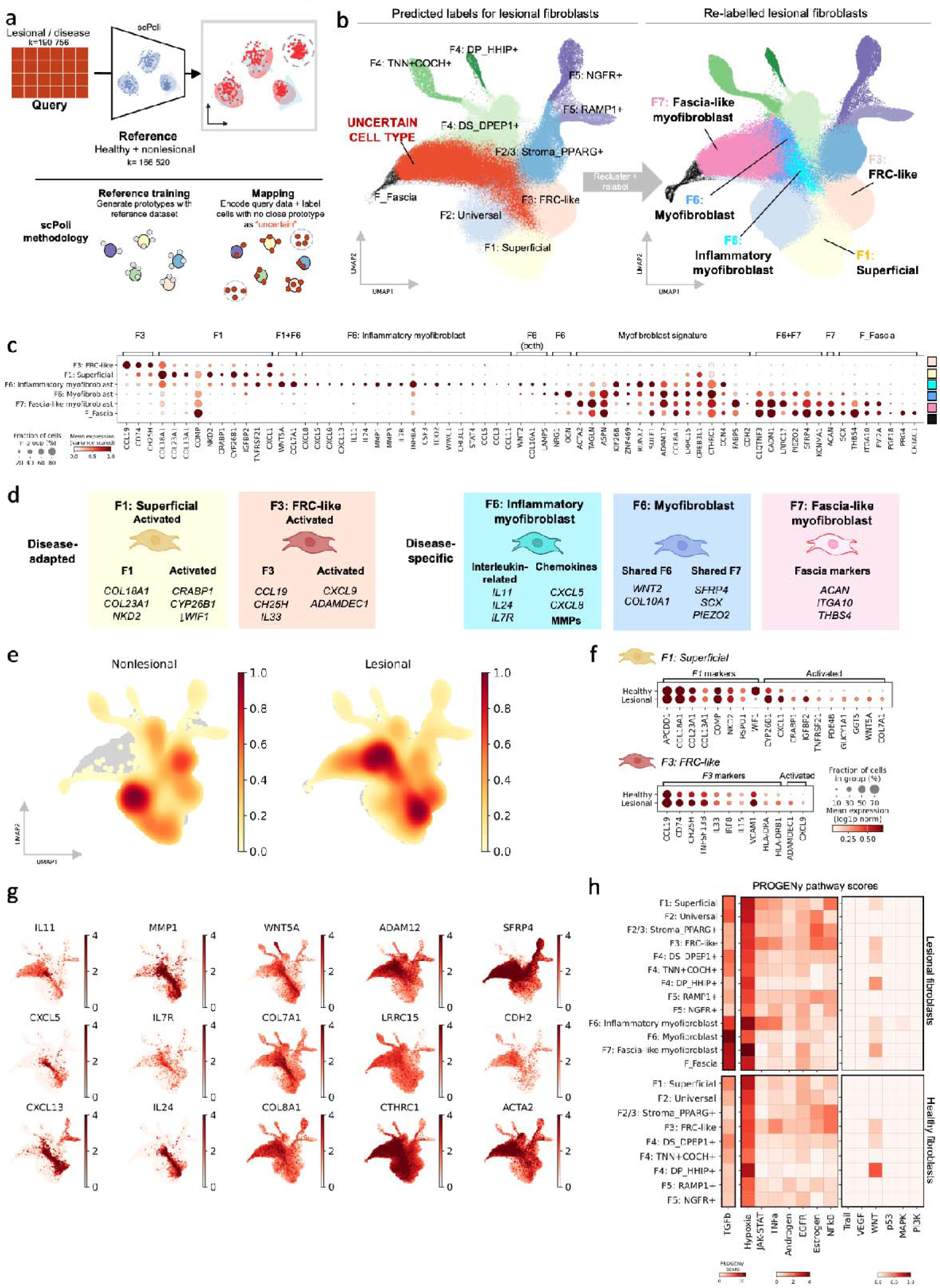
Prototype meta-learning to identify disease-adapted and disease-specific populations. a) Overview of reference-mapping approach used for integration, where lesional/diseased data was mapped using a pre-trained model. scPoli methodology for reference-mapping specifically, using meta prototypical learning, is shown under the line. b) UMAP of embedding from scPoli with predicted cell type labels and relabelled populations after clustering. c) Dot plot of marker gene expression for disease-adapted and disease-specific fibroblast populations See Extended Data Fig 6a for differentially expressed genes for disease fibroblast populations. d) Summary of disease-adapted and disease-specific populations. e) Density of cells in embedding by site status. f) Gene expression in *F1* and *F3* fibroblasts from health and disease, including differentially expressed genes in lesional/diseased states. g) UMAP visualisation of genes associated with myofibroblast subtypes. h) PROGENy pathway scores for fibroblasts from lesional and healthy samples.

scPoli mapped 121,167 diseased cells to existing labels (*F1-F5*) from the healthy/non-lesional reference. The same marker genes used for the healthy/non-lesional reference clearly delineated *F1-F5* fibroblast subtypes in the new disease data (Extended Data Fig. 5a), although some marker genes lost specificity as they were upregulated in novel disease subtypes (e.g. *ASPN*, *COL11A1*, *CTHRC1*, and *COMP*) (Extended Data Fig. 5b). scPoli classified 69,589 fibroblasts from the disease data as uncertain. These unmapped disease-related fibroblasts were manually annotated to reveal two disease-adapted and two disease-specific fibroblast subtypes (Fig. 3b-d and Extended Data Fig. 6a).

#### Disease-adapted fibroblast subtypes

Disease-adapted fibroblast subtypes marked as uncertain by scPoli (Extended Data Fig. 5c) resembled a healthy fibroblast counterpart but expressed genes with additional molecular properties and were often expanded in disease settings (Fig. 3e and Extended Data Fig. 5d). The first disease-adapted fibroblasts resembled *F1: Superficial* fibroblasts in healthy skin (*COL18A1*, *COL23A1*, *NKD1*) (Fig. 3f). In disease, this population upregulated genes suggestive of regenerative function (*CRABP1* and *CYP26B1)* and downregulated the Wnt antagonist *WIF1* (Fig. 3f). *CRABP1* and *CYP26B1* are markers of superficial/upper wound fibroblasts in mice^52,68^, which are thought to be the source of wound-induced hair follicle neogenesis in mice^69^. *Crabp1*+ fibroblasts also are associated with the regenerative potential of reindeer velvet skin^70^ and early-gestational human prenatal skin^71^. CRABP1 and CYP26B1 modulate intracellular retinoic acid levels^72^, suggesting a similar retinoic acid gradient in human skin to that observed in mice, which regulates early myofibroblast differentiation^52^.

The second disease-adapted fibroblast subpopulation was *F3: FRC-like,* which expressed higher amounts of *CXCL9* and *ADAMDEC1* (Fig. 3f). *CXCL9* has been reported as an activation marker of FRCs in lymph nodes and Peyer’s patches^53^, and in a mouse colitis model, *Adamdec1* was involved in mucosal matrix remodelling and healing^73^. *CXCL9*+ and *ADAMDEC1*+ fibroblasts likely represent activated *F3: FRC-like* populations, as suggested by previous work^6^. *CCL19*+*ADAMDEC1*+ fibroblasts have been reported as a gastrointestinal tract-specific population^6^, but we identify that these fibroblasts are also present in human skin.

#### Disease-specific fibroblast subtypes

Disease-specific fibroblasts were absent in healthy skin and only observed in disease. There were three disease-specific subtypes: *F6: Inflammatory myofibroblasts*, *F6: Myofibroblasts,* and *F7: Fascia-like myofibroblasts.* These subtypes were characterised by elevated extracellular matrix (ECM) expression (*COL3A1*, *COL5A1, COL8A1, POSTN*, *TNC*, *SPARC, CTHRC1*), increased contractility markers (*ACTA2*),^74^ and expression of other myofibroblast-associated genes (*LRRC15*, *ADAM12*) (Fig. 3c,g and Extended Data Fig. 5e)^5,37,75–78^.

*F6: Inflammatory myofibroblasts* expressed genes encoding chemokines (*CXCL5*, *CXCL6*, *CXCL8*, *CXCL13*), interleukins (*IL11, IL24*), and matrix metalloproteinases (*MMP1*, *MMP3*) (Fig. 3c,d). *F6: Inflammatory myofibroblasts* also showed higher *CCL19* expression than other myofibroblast populations (Fig. 3c). *CXCL5*, *CXCL6*, and *CXCL8* are chemoattractants primarily for neutrophils, which characterise the early inflammatory phase of wound healing^79^. *IL11* is a described driver of fibrosis and cancer pathogenesis in mouse models^80–83^.

*F6: Myofibroblasts* and *F7: Fascia-like myofibroblasts* were distinguished by high expression of genes such as *SFRP4, LRRC17,* and *PIEZO2*, as well as distinct expression of genes encoding cellular retinoic-acid binding proteins (*FABP5*^hi^*CRABP1*^lo^) (Fig. 3c). Expression of ECM and contractility genes was also typically highest in *F6: Myofibroblasts* and *F7: Fascia-like myofibroblasts* (Extended Data Fig. 5e). *PIEZO2* encodes the major mechanotransducer of mammalian proprioceptors^84^, and mechanical force can activate myofibroblasts and drive fibrosis^85^. *F7: Fascia-like myofibroblasts* were distinguished from *F6: Myofibroblasts* by their higher expression of genes we previously observed in fascial fibroblasts (*ITGA10*, *THBS4*, *PDLIM3*). *CDH2* showed a modest increased expression in *F7: Fascia-like myofibroblasts* (Fig. 3c). CDH2 mediates the swarm-like migration of fascial fibroblasts into the dermis in mice^50^.

Finally, we used PROGENy pathway analysis to further understand the signalling pathways that are altered in disease-adapted and disease-specific fibroblast populations. Wnt signalling was elevated in disease-adapted *F1* and all disease-specific populations (*F6-F7*) (Fig. 3h). JAK-STAT signalling was elevated in *F3: FRC-like* and *F6: Inflammatory myofibroblasts*. Hypoxic signalling, which regulates early myofibroblast differentiation in mice^52^, was elevated in *F6: Inflammatory myofibroblasts* and *F7: Fascia-like myofibroblasts*. TGFβ signalling, which is the main controller of the myofibroblast state^2^, was highest in *F6: Myofibroblasts* and *F7: Fascia-like myofibroblasts* (Fig. 3h). This analysis suggests that human skin disease-adapted and disease-specific fibroblast subtypes upregulate specific signalling pathways in addition to TGFβ signalling that can be potentially targeted for therapy, including retinoic acid activity, hypoxic signalling, and JAK-STAT signalling.

### Fibroblast compositional signatures characterise the stroma of distinct skin diseases

We next hypothesised that specific pathogenic processes in tissue can be distinguished by the fibroblast composition of the stroma. To test this, we first grouped the 23 skin diseases into predominantly ‘inflammatory’, ‘cancer-related’, or ‘scarring/fibrotic’ categories (Fig. 4a). Within diseases that were predominantly inflammatory, we further distinguished them by clinically-determined risk of scarring (Methods). We considered neurofibroma as a distinct entity as it was the only case of benign neoplasia. Using this approach, we demonstrated that different disease conditions are characterised by distinct fibroblast compositions (Fig. 4a,b).

**Figure 4.**
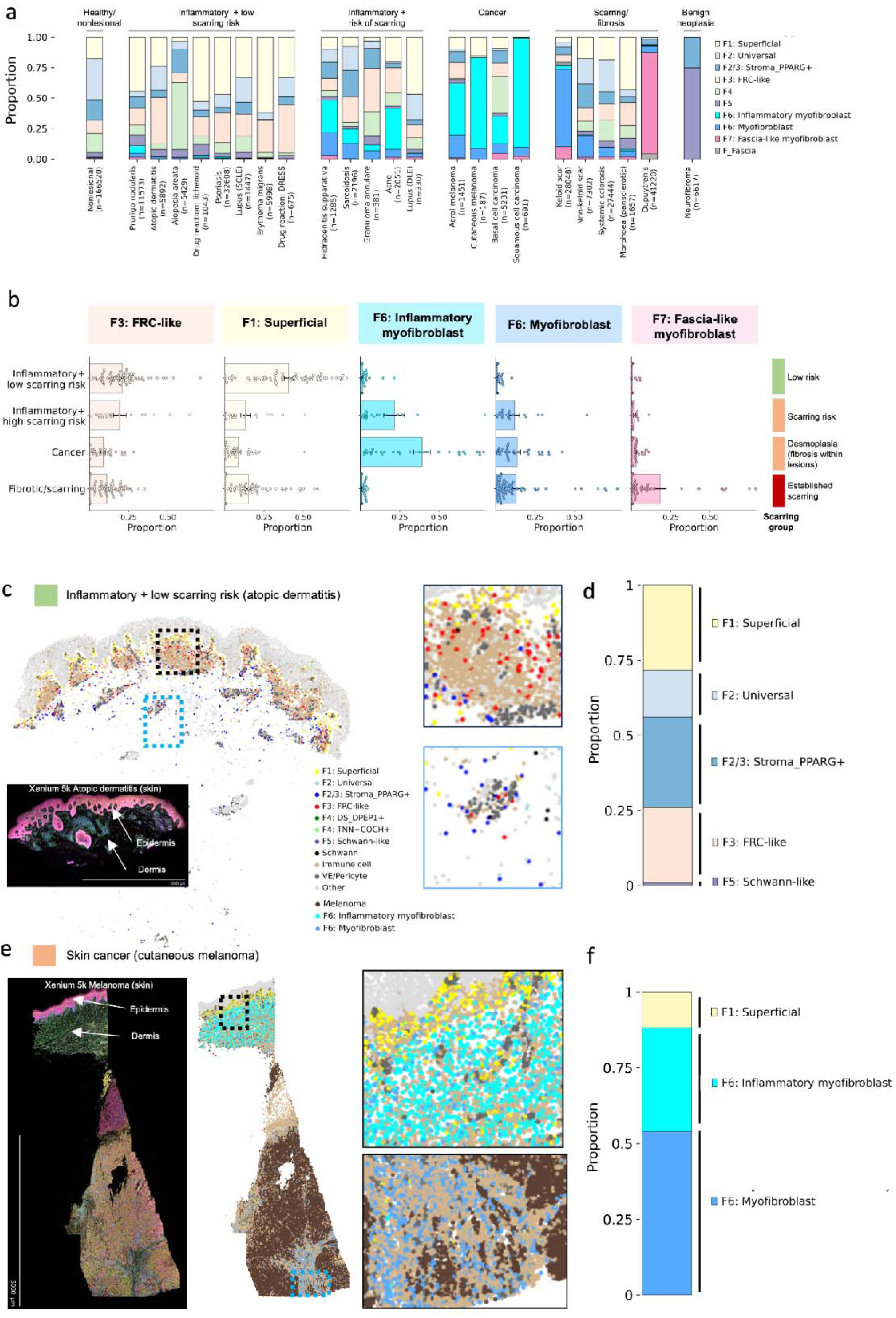
Fibroblast compositional signatures characterise the stroma of distinct skin diseases. a) Proportion of fibroblast populations by individual disease. b) Proportion of disease-adapted and disease-specific fibroblast subtypes by disease category (mean +-standard error of the mean). Scarring risk group was based on clinical profiles of these diseases (see Methods). c) Xenium 5k data for lesional/inflamed atopic dermatitis skin, with cells coloured by cell type. d) Proportion of fibroblast subtypes in lesional/inflamed atopic dermatitis skin. e) Xenium 5k data for cutaneous melanoma, with cells coloured by cell type. f) Proportion of fibroblast subtypes in cutaneous melanoma.

‘Inflammatory low-scarring risk’ skin disease stroma was characterised by a high prevalence of *F1: Superficial* fibroblasts, consistent with their regenerative-associated gene profile, and *F3: FRC-like* fibroblasts (Fig. 4b). *F6-F7* myofibroblasts were rarely present. ‘Inflammatory high-scarring risk’ stroma and cancer-related stroma were characterised by a uniquely high prevalence of *F6: Inflammatory myofibroblasts* (Fig. 4b). *F6: Myofibroblasts* were also elevated, with a similar prevalence to that in established fibrosis/scarring skin disorders. *F6: Inflammatory myofibroblasts* were transcriptionally enriched with genes reported for inflammatory cancer-associated fibroblast (iCAFs) in skin cancer, whereas *F6: Myofibroblasts* showed a matrix-producing phenotype (*COL1A1*, *COL3A1*, *POSTN*, *LRRC15*) that characterises myofibroblastic CAFs (myoCAF) (Extended Data Fig. 7a)^86^. Heterogeneity of iCAFs is recognised^87^, which we also observed in skin cancer (Extended Data Fig. 7b).

Established fibrotic disorders, including conventional scars, were characterised by a high prevalence of *F6: Myofibroblasts* and *F7: Fascia-like myofibroblasts*, without significant *F6: Inflammatory myofibroblast* populations (Fig. 4b). Fibroproliferative disorders (keloid scarring and Dupuytren contracture), in which scars actively expand in size, showed a very high prevalence of *F6: Myofibroblasts* and *F7: Fascia-like myofibroblasts* (Fig. 4a).

We were able to distinguish and characterise the fibroblasts comprising neurofibromas, benign tumours of peripheral nerve-sheath including in the skin, using our comprehensive skin fibroblast atlas.^10^. We identified that neurofibroma fibroblasts consist primarily of *F5: Schwann-like* and *F2/3: Stroma_PPARG+* fibroblasts (Fig. 4a), and validated the gene expression for these populations (Extended Data Fig. 7c). This finding also supports a nerve-associated location for *F5: Schwann-like* fibroblasts, which we observed by spatial transcriptomic analysis (Extended Data Fig. 3a,f and 4h). Overall, these data point towards *F6: Myofibroblasts* and *F7: Fascia-like myofibroblasts* as pro-fibrotic populations.

To validate the distinct disease stromal profiles and investigate spatial architecture, we performed spatial genomics analysis (Xenium 5k-plex) of inflamed atopic dermatitis skin (low-scarring risk stroma) and interrogated a publicly-available cutaneous melanoma Xenium 5k-plex dataset (Fig. 4c-f and Extended Data Fig. 7d-g).^88^ Stromal composition profiles by Xenium analysis were in accordance with scRNA-seq data (Fig. 4a-b,d-f). Low-scarring risk stroma comprised *F1-F5* fibroblasts, without significant myofibroblasts. In cancer, aside from *F1,* the entire stroma comprised *F6: Inflammatory myofibroblasts* and *F6: Myofibroblasts* (Fig. 4e,f), which correspond to iCAFs and myoCAFs, respectively (Extended Data Fig. 7a)^86,89^. Spatially, in low-scarring risk stroma, *F3: FRC-like* cells were enriched in the superficial perivascular niche (Fig. 4c). In cancer, *F6: Inflammatory myofibroblasts* and *F6: Myofibroblasts* showed a non-exclusive enrichment in superficial and deep skin, respectively. In both diseases, *F1: Superficial* fibroblasts predominantly localised to the epidermis, in keeping with continued maintenance of skin epithelium during disease.

The dramatic shift in stromal composition in cutaneous melanoma likely contributes to novel, disease-specific immune niches. For example, *CXCL13*, a known B cell chemoattractant, was expressed by *F6: Inflammatory myofibroblasts* (Extended Data Fig. 7f) but not significantly expressed in healthy skin fibroblasts (Extended Data Fig. 7g). In keeping with this, B cells, which are rare in healthy skin, were prevalent in cutaneous melanoma, particularly in tissue regions containing *F6: Inflammatory myofibroblasts* (Extended Data Fig. 7e,f). This lends support to disease-specific fibroblast phenotypes modulating local immune niches.

### Origin of skin disease-specific myofibroblasts

The origin of disease-specific myofibroblasts that contribute to fibrosis is a key question with significant clinical relevance, but remains poorly understood in human tissues. Fibroblasts are tissue resident, and thus intermediate states of myofibroblast differentiation are likely to be captured in the molecular snapshots of skin diseases within the datasets we analysed. We therefore performed trajectory analysis of fibroblasts in disease states using four different computational approaches to gain further insights into the disease-specific fibroblast (myofibroblast subtypes) differentiation in the skin (Fig. 5a-c and Extended Data Fig. 8a-c).

**Figure 5.**
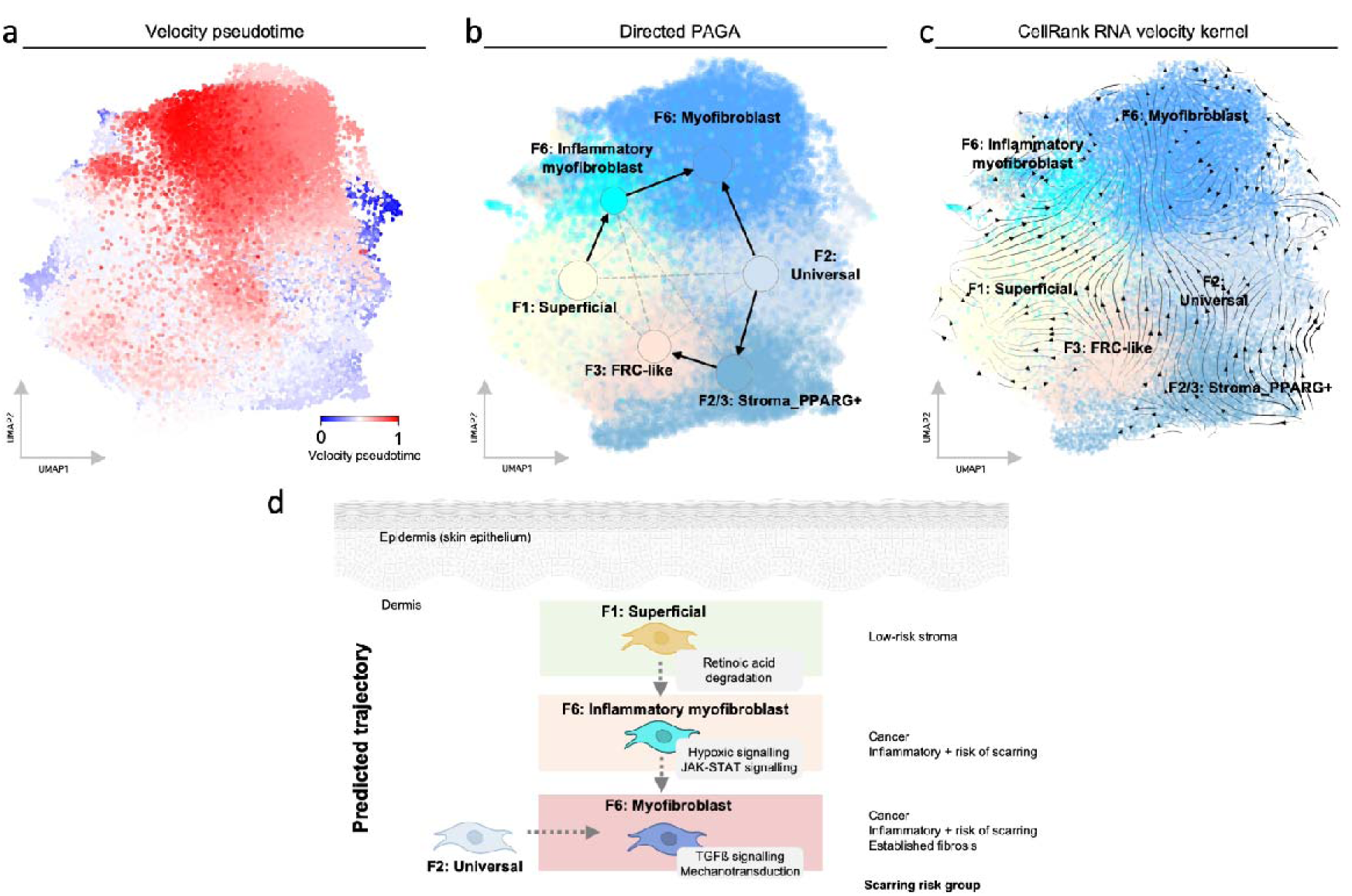
Origin of skin disease-specific fibroblast subtypes, a) Velocity pseudotime, b) Partition-based graph abstraction (PAGA) overlaid on UMAP, with direction calculated from RNA velocity, and c) Velocity kernel from CellRank2 for lesional fibroblasts. For further details see Methods. d) Schematic of predicted trajectories. Dotted arrows indicate predictions with multiple lines of evidence. Fibroblast populations are coloured by the predominant scarring/fibrosis risk observed in an earlier analysis: green (prevalent in low risk scarring stroma), orange (prevalent in scarring risk stroma and cancer), red (prevalent in established scarring/fibrotic disorders). Grey boxes indicate signalling pathways identified in our gene expression/pathway analysis.

RNA velocity (scVelo, CellRank2 Velocity kernel) and graph-based approaches (PAGA, Monocle 3) revealed a consistent inferred trajectory for *F6: Myofibroblast* as a terminally-differentiated myofibroblast state, arising from either *F2: Universal* fibroblasts directly or from *F1: Superficial* fibroblasts via an intermediate *F6: Inflammatory myofibroblast* state (Fig. 5a-d and Extended Data Fig. 8b-c). While these predictions require validation, our inferred trajectories (Fig. 5d) are consistent with *in vivo* lineage tracing studies in mice that indicate an intermediate inflammatory myofibroblast state^52,90^, as well as a direct trajectory from *F2: Universal* (reticular) fibroblasts to myofibroblasts^56^.

### Human cross-tissue disease fibroblast populations

Previous studies have analysed fibroblasts from multiple diseases across human tissues and defined their marker genes, but the fibroblast populations were named differently in each study^4–6^. We therefore assessed the expression of these disease-related fibroblasts marker genes in our skin fibroblast populations (Fig. 6a and Extended Data Fig. 9a) and derived a consensus nomenclature to link these fibroblast populations across human tissues.

**Figure 6.**
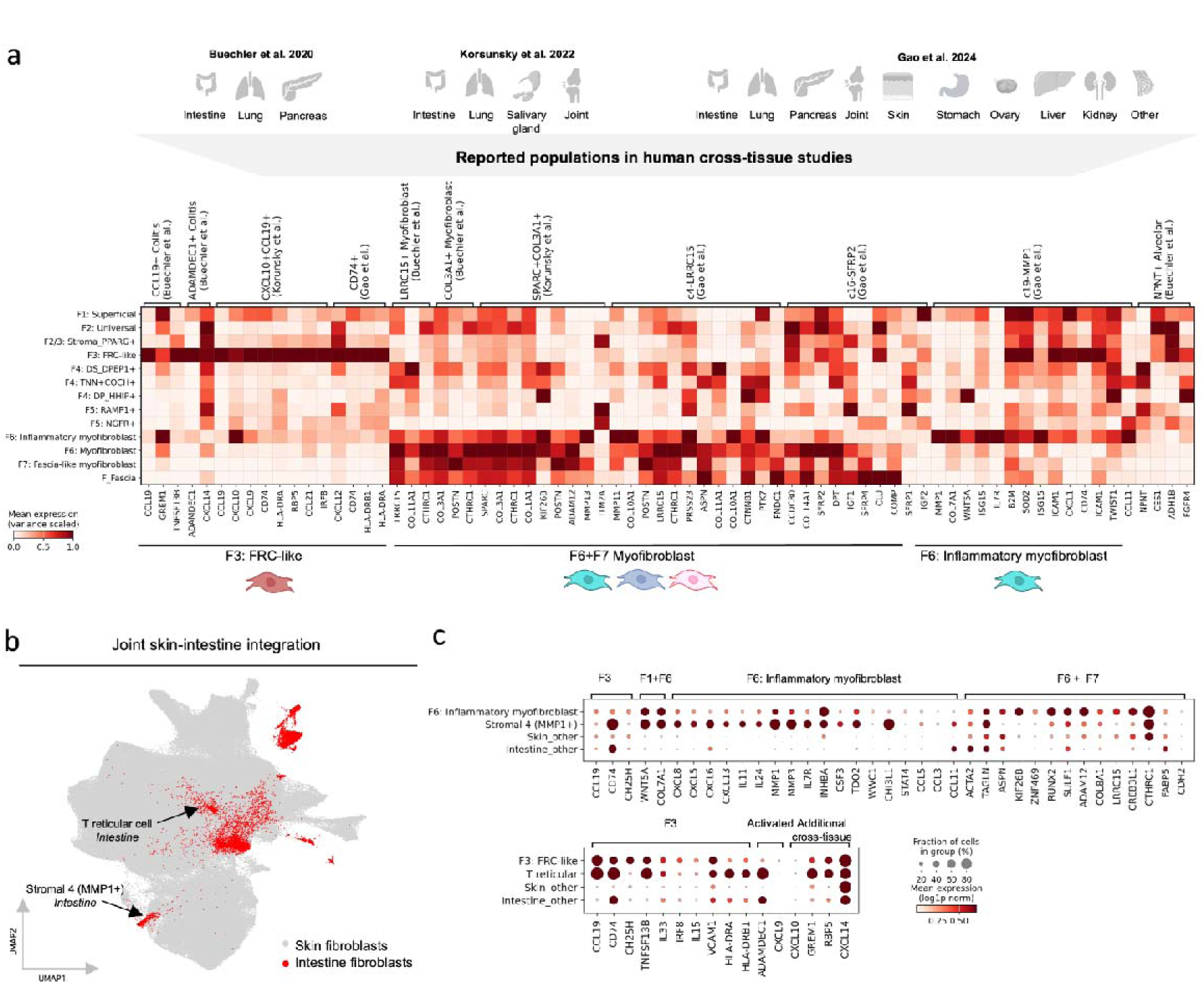
Human cross-tissue disease fibroblast populations. a) Human tissues used in cross-tissue fibroblast studies and heatmap of gene expression of marker genes for reported cross-tissue fibroblast populations in our skin fibroblast populations. b) UMAP visualization of scVI embedding for joint human skin-intestine fibroblast integration. c) Dotplot of marker genes for *F6: Inflammatory myofibroblasts* (from skin) in *F6: Inflammatory myofibroblasts* (skin) and *Stromal 4 (MMP1+)* (intestine) populations; dotplot of marker genes for *F3: FRC-like* (from skin) in *F3: FRC-like* (skin) and *T reticular cells (*intestine) populations (same scale for both plots).

All cross-tissue studies reported a population with similarity to *CCL19*+ *FRC-like* fibroblasts with a shared gene expression signature (*CCL19*, *CD74*, *HLA-DRA*) (Fig. 6a). This population is expanded in rheumatoid arthritis and IBD^6^ and observed in cancers in multiple tissues^5^. Myofibroblasts reported across tissues were similar to skin *F6-F7* myofibroblasts broadly (*COL3A1*, *CTHRC1*, *ADAM12*, *LRRC15*)*. F6: Inflammatory myofibroblasts* showed specific similarity to an *MMP1+* population (Fig. 6a), which was recently identified in a large-scale integration of predominantly cancer fibroblasts and linked to immunotherapy response^5^. *F1: Superficial* fibroblasts did not correlate with fibroblasts in other human tissues, which may reflect distinct functional requirements across epithelial barrier tissues.

To more comprehensively investigate if skin disease-adapted and disease-specific fibroblasts are observed in another tissue directly, we next jointly integrated our skin fibroblast data with independent human intestinal scRNA-seq data from healthy and IBD intestinal tissue (Fig. 6b and Extended Data Fig. 9b)^91^. This approach uses the whole transcriptome profile instead of restricted marker genes. The joint integration confirmed that *F3: FRC-like* fibroblasts are analogous to reported cross-tissue *CCL19*+ populations. *F3: FRC-like* cells (skin) clustered with *T zone reticular* cells (intestine) (Extended Data Fig. 9b), and both skin and intestinal populations highly expressed FRC-related genes (*CCL19*, *CD74*, *TNFSF13B*, *VCAM1*, *HLA-DRA, IL33*) (Fig. 6c). This result suggests a conserved immune-interacting population that resembles FRCs across human organs, including non-lymphoid tissue.

Interestingly, skin *F6: Inflammatory myofibroblasts* and intestinal *Stromal MMP1+* fibroblasts clustered closely (Extended Data Fig. 9b) and shared similar gene expression (Fig. 6c). Intestinal *Stromal MMP1+* fibroblasts expressed the same gene signature as cross-tissue *MMP1+* fibroblasts (*MMP1, COL7A1, WNT5A*, *IL7R, IL11*)^5^ (Extended Data Fig. 9c) and *inflammatory-associated fibroblasts* (*IL11*+*IL24*+) (Extended Data Fig. 9d,e), which are associated with non-response to biologic therapy in IBD^64^. These results suggest that inflammatory myofibroblasts (*MMP1, COL7A1, WNT5A*, *IL7R, IL11*) are a cross-tissue fibroblast population present in inflammatory disorders and cancer across multiple human tissues^5^.

### Adult skin F3: FRC-like fibroblasts regulate immune niches and potentially arise from prenatal skin LTo cells

In the developing Peyer’s patch, a secondary lymphoid organ of the intestine, FRCs arise from both endothelial LTo cells and *CCL19*+ mesenchymal LTo cells^92^. We previously reported *CCL19+* fibroblasts in prenatal skin^71^ with a similar gene expression profile to adult skin *F3: FRC-like* fibroblasts and intestinal *T zone reticular* fibroblasts. We therefore hypothesised that prenatal skin *CCL19*+ fibroblasts are similar to intestinal mesenchymal LTos and potentially give rise to adult skin *F3: FRC-like* cells in a parallel manner to the intestine.

To investigate this, we integrated publicly-available human prenatal intestinal fibroblasts (147 628 cells) with human prenatal skin fibroblasts (269 770 cells)^71,91^. Strikingly, prenatal skin *CCL19*+ cells and intestinal mesenchymal LTo cells clustered together (Fig. 7a). Prenatal skin *CCL19*+ cells expressed known mesenchymal LTo markers^91^, including *CCL19*, *CCL21*, *CXCL13*, *MADCAM1*, *FDCSP*, and *TNFSF11* (RANKL) (Fig. 7b). This suggests that LTo-like *CCL19+* cells in prenatal skin may differentiate into adult skin *F3: FRC-like* cells, similar to intestinal tissue. The skin does not harbour the equivalent of Peyer’s patch, and thus the role of LTo-like cells during development beyond the formation of secondary lymphoid organs remains to be explored.

**Figure 7.**
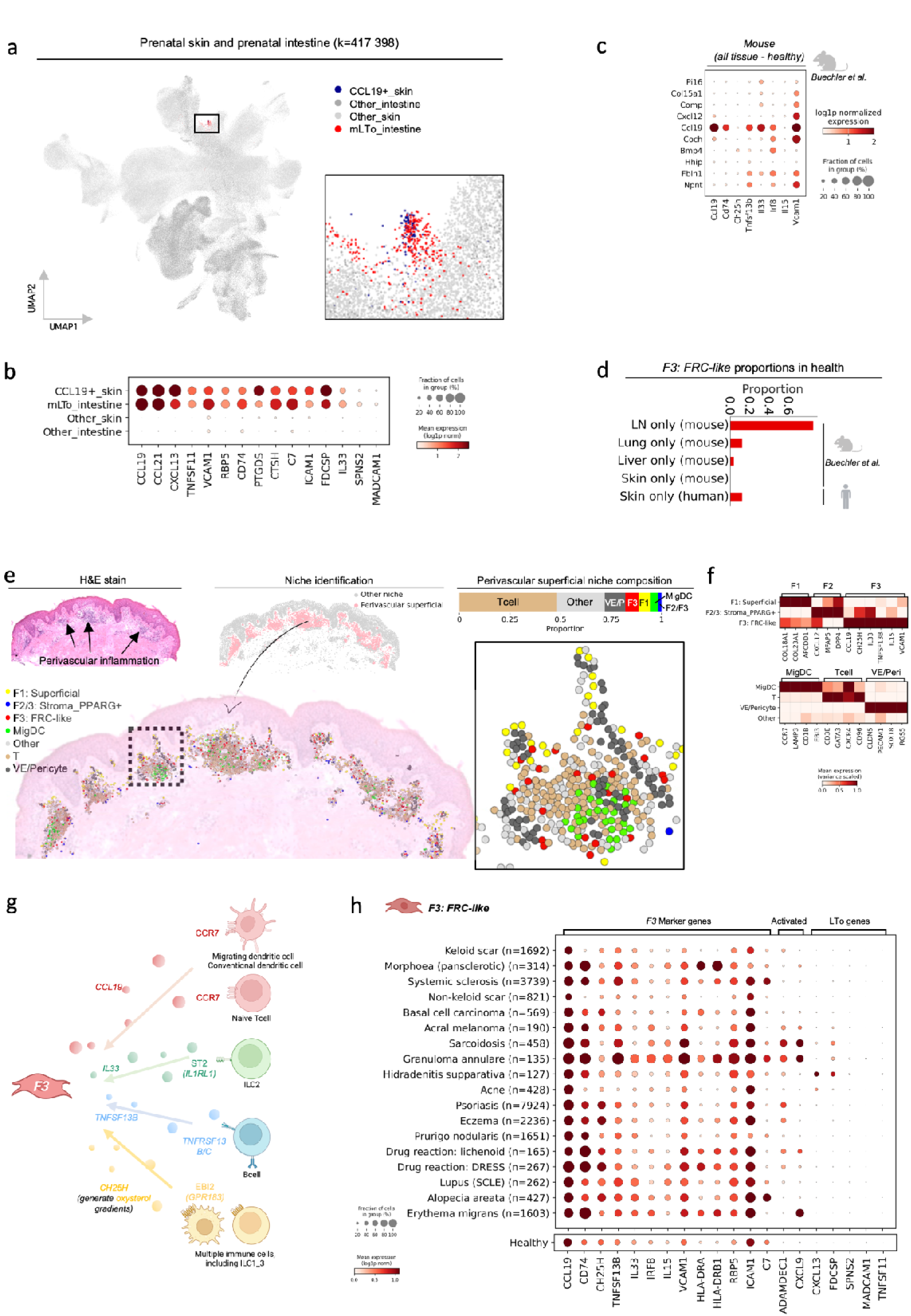
Adult skin *F3: FRC-like* fibroblasts regulate immune niches and potentially arise from prenatal skin LTo cells. a) UMAP visualisation of scVI embedding for prenatal human skin and intestine. Insert: Cluster of LTo-like cells. (m)LTo: (mesenchymal) lymphoid tissue organizer. b) Dotplot of marker gene expression for LTo-like cells. c) Dotplot of expression of marker genes for human skin *F3: FRC-like* fibroblasts in mouse Ccl19+ fibroblasts from the mouse steady-state cross-tissue atlas (Buechler et al). d) Proportion of *Ccl19+* fibroblasts (labels from original study) in mouse in different healthy tissues and of *F3: FRC-like* fibroblasts in healthy skin. e) H&E slide of lesional atopic dermatitis skin with annotation of perivascular infiltrate regions. Niche identification and proportion of cells in the perivascular superficial niche. Composition of the perivascular superficial niche in 10x Genomics Xenium. Insert: zoomed in version of perivascular niche cluster. H&E: Haematoxylin and eosin f) Heatmap of marker gene expression for fibroblast and non-fibroblast populations in atopic dermatitis superficial perivascular niche. g) Schematic representation of cell-cell interactions for *F3: FRC-like* fibroblasts from CellPhoneDB and previous results in type 2 immunity (Dahlgren et al.). See Methods for more detail. h) Dotplot of gene expression for *F3: FRC-like* fibroblasts from healthy skin and from skin across different diseases (lesional), for diseases with a minimum of 100 *F3: FRC-like* fibroblasts.

As no LTo-like fibroblasts have been identified in mouse embryonic skin^93^, we asked whether *F3: FRC-like* fibroblasts were unique to human skin. We used the cross-tissue mouse fibroblast atlas data to perform cross-species analysis^4^. We found that *F3: FRC-like* cells are the equivalent of mouse *Ccl19*+ fibroblasts, with similar expression of homologous genes (Fig. 7c). Using labels from the original mouse fibroblast atlas, we confirmed that *Ccl19*+fibroblasts were predominantly a lymphoid organ population in mouse, but also present in non-lymphoid tissues in which *Ccl19*+ fibroblasts have been reported, including lung and liver (Fig 7d)^94^. *Ccl19*+ fibroblasts were very rare in healthy mouse (flank) skin (Fig 7d). In contrast, *F3: FRC-like* fibroblasts were comparatively common in human adult skin (Fig. 7d), including across different anatomical sites (Extended Data Fig. 1e).

Next, we further explored the role of *F3: FRC-like* fibroblasts in atopic dermatitis. Expanded perivascular inflammatory infiltrate is characteristic of atopic dermatitis lesional skin (Fig. 7e), and we had previously observed that *F3: FRC-like* cells were associated with the superficial immune cell niche (Fig. 4c and Extended Data Fig. 3a,c-d). To quantify this association, we used NicheCompass^95^ to identify and deconvolve the perivascular immune niche, identifying *F3: FRC-like* as the most common fibroblast subtype (Fig. 7e). We then further annotated immune cells in this niche and identified *CCR7*+*LAMP3*+ migrating dendritic cells (MigDCs) and T cells (Fig. 7e,f)^31^. Using receptor–ligand mapping^96^, we predicted potential interactions of *F3: FRC-like* cells with dendritic cells (CCL19-CCR7) and T cell subsets (CXCL12-CXCR4, PLA2G2A-very late Ag-4(VLA4; integrin α4β1), VCAM1-VLA4) (Fig. 7g and Extended Data Fig. 10a). These results suggest that *F3: FRC-like* fibroblasts maintain the expanded perivascular inflammatory infiltrate niche and facilitate DC and T cell interactions, analogous to the role of *T reticular cells* in lymphoid organs. Importantly, such perivascular inflammatory structures are not typically observed in mouse skin^54^.

The perivascular inflammatory structures observed in many human inflammatory skin diseases are poorly organized and thus not typically considered to be tertiary lymphoid structures (TLS)^54^. However, TLS is reported in human skin in hidradenitis suppurativa and cutaneous T-cell lymphoma^97,98^. Given the role of stromal cells in TLS formation^99^, we investigated the gene expression profile of *F3: FRC-like* fibroblasts across different skin diseases. We observed that LTo-associated genes, such as *CXCL13* and *FDCSP*, were not expressed in *F3: FRC-like* cells in healthy skin, but were upregulated by both *F3: FRC-like* and *F6* myofibroblast populations in select diseases, including hidradenitis suppurativa (Fig. 7h and Extended Data Fig. 9b). *CXCL13* appears to be an important chemokine for TLS formation^97,98,100^, and its expression in select diseases may distinguish perivascular inflammatory infiltrates from TLS. Overall, our findings suggest that skin *F3: FRC-like* cells are a cross-tissue human fibroblast population with an important role maintaining the superficial perivascular immune cell niche. *F3: FRC-like* fibroblasts also potentially mediate TLS formation and B-cell homing in select diseases through re-activation of a developmental gene program.

## Discussion

In this study, we defined 6 major fibroblast clusters and their spatial niches in healthy skin, identified three novel myofibroblast states in skin disease, and described the fibroblast subtype composition within the stroma across different skin diseases. Previous skin fibroblast data integration efforts have not included disease categories, single-cell resolution spatial transcriptomic data, nor linked populations to cross-tissue fibroblast states^49,101^. Through including these factors, we harmonised skin fibroblast subtype nomenclature in health and disease, spatially resolved distinct fibroblast anatomical niches, and identified conserved fibroblast subtypes in human diseases affecting multiple tissues^5^.

An immunomodulatory role for fibroblasts is best exemplified by FRCs in lymphoid organs^102^. We identified that *F3: FRC-like* fibroblasts were an important cell type in the superficial immune cell niche in human skin, but were rare in mouse skin, in keeping with previous murine studies^54,103^. We also identified LTo-like cells in prenatal human skin, which have not been identified in prenatal mouse skin^93^. These observations suggest an important difference in fibroblast composition, and likely origin, between human and mouse skin with respect to their contribution to the cutaneous immune niche. Accordingly, although TLS-like structures have been proposed in mouse skin, such as inducible skin[associated lymphoid tissue (iSALT) in a contact dermatitis model^104^, these predominantly comprise DC and T-cells, without FRC-like stromal cells^105,106^. An absence of *F3: FRC-like* fibroblasts in mouse skin would elegantly explain both why fibroblasts are not reported in mouse iSALT and why the prominent perivascular infiltrate structures that characterise many human inflammatory skin diseases are not reported in mouse^54^. In agreement with a species difference, iSALT in human skin in secondary syphilis is associated with CXCL13+ fibroblast-like cells in the superficial dermis^107^. The identity of this fibroblast cell was not clear^107^, but our results suggest *F3: FRC-like* and/or *F6: Inflammatory myofibroblast* as likely candidates. Overall, our results suggest cross-species differences in how dermal fibroblasts mediate immune cell niches and how they respond to inflammation, reflecting known and emerging differences during tissue morphogenesis and maturation between human and mouse skin and lymphoid tissue^71,102^.

Recently, tissue-specific fibroblasts that support alveolar epithelium in mouse lung have been reported to give rise to fibrotic myofibroblasts via an inflammatory intermediate.^90^ *Universal* fibroblasts, which are found across multiple human tissues, have also been suggested as a source of myofibroblasts^4^. This led to a recent proposal that there are two potential sources for myofibroblasts: a tissue-specific population and a *Universal* population^108^. Our results support this proposal, with evidence for two trajectories: one from *F1: Superficial* (epithelium-associated) via an *F6: Inflammatory myofibroblast* intermediate, and one from *F2: Universal* via direct differentiation. Our results raise the possibility that the differentiation of epithelium-related, or *Superficial*, fibroblasts to myofibroblasts is not unique to lung tissue, but may also be observed in other tissues, including skin. Further understanding the steps of myofibroblast differentiation in human tissue is a vital step towards developing highly efficacious anti-fibrotic treatments. The advent of single-cell resolution spatial transcriptomics, together with the ability to temporally sample human skin, provides a promising avenue to further decode myofibroblast differentiation.

Drug modulation of fibroblast states has great clinical potential due to their key roles in wound healing; fibrosis; and modulation of immune cell infiltrate^76,109^. We identify that *F3: FRC-like* and *F6: Inflammatory myofibroblasts* represent interesting target populations as they are present in different organs across multiple diseases, including inflammatory disease and cancer. Inhibition of terminally-differentiated myofibroblasts that characterise fibrosis also presents an attractive therapeutic avenue, which in mice can be achieved through HIF1α inhibition and JAK inhibition^52,109^.Whether the distinct patterns of fibroblast composition across disease categories is reflected in distinct and therapeutically-targetable ECM profiles warrants further investigation.

Our integrated skin fibroblast dataset is freely available for viewing and download at (to be added upon publication). Our fibroblast annotation provides a basis for future studies and is available for interrogation and feedback (https://celltype.info/project/388). In summary, our study provides a foundational resource for fibroblast transcriptomic states in health and across distinct disease categories in skin tissue, including cancer, fibrosis, and immune-mediated conditions.

**Extended Data Fig. 1.**
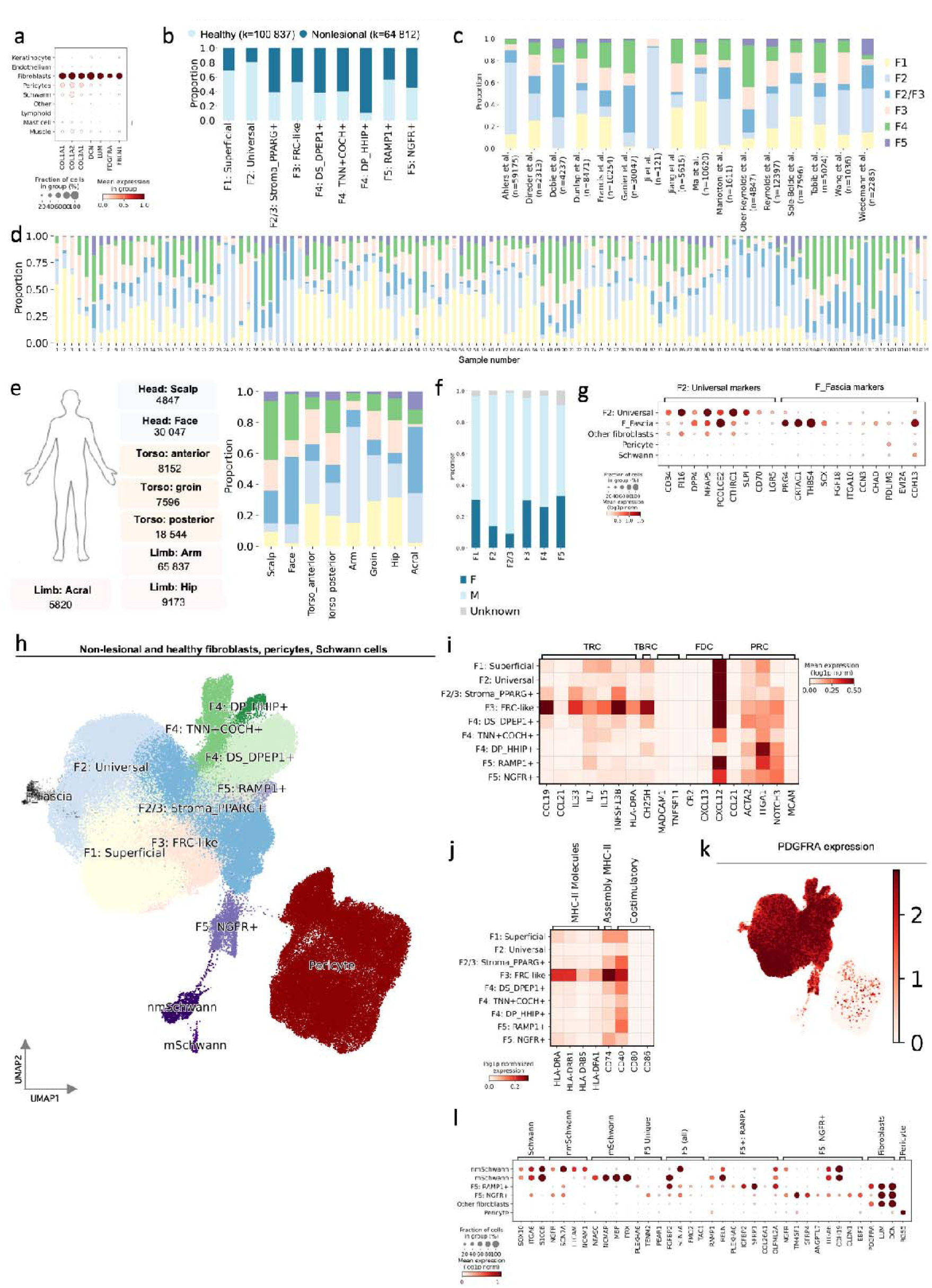
Metadata and integration with Schwann cells, pericytes, and fascial cells. a) Marker genes for fibroblast selection. b) Proportion of fibroblasts by site status, c) dataset, d) sample, e) anatomical location, and f) sex. g) Dot plot of marker genes for fascial fibroblasts. h) UMAP of fibroblast integration including Schwann cells, fascial fibroblasts, and pericytes. i) Heatmap of reported marker genes for fibroblastic reticular cell (FRC) subsets. TRC: T reticular cell. TRBC: T-B border reticular cells. FDC: Follicular dendritic cell. PRC: Perivascular reticular cell. j) Heatmap of MHC-II and related molecules. k) UMAP visualisation of additional fibroblast integration showing *PDGFRA* expression. l) Dotplot of Schwann cell and *F5: Schwann-like* fibroblast gene expression.

**Extended Data Fig. 2.**
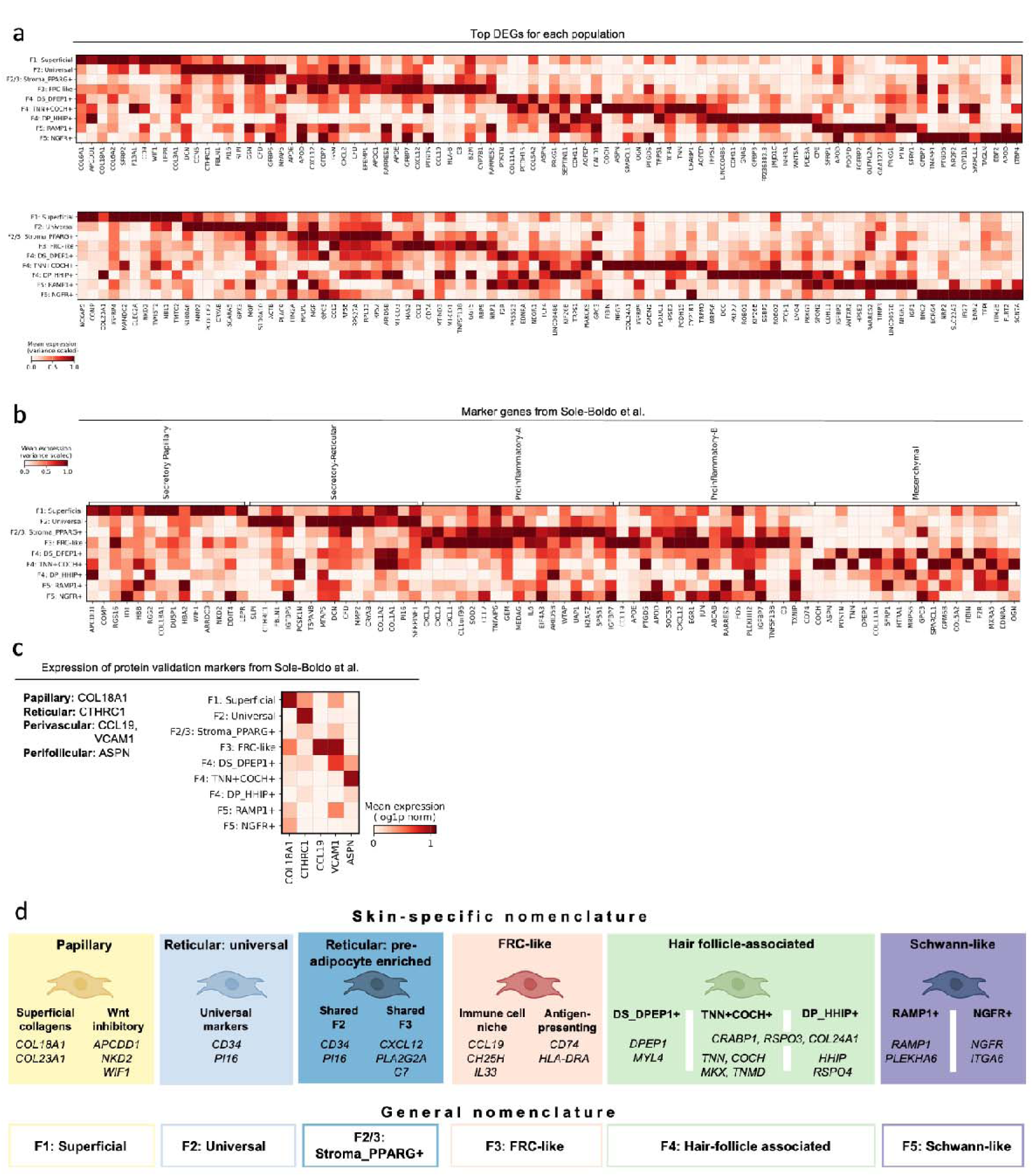
Differentially expressed genes for healthy fibroblasts and skin-specific nomenclature. a) Heatmap of top differentially expressed genes (DEGs) for each fibroblast population. Top row: Top 10 DEGs. Bottom row: 11-20 top DEGs. b) Heatmap of expression of marker genes reported by Sole-Boldo et al. c) Heatmap of expression of genes used for protein validation by Sole-Boldo et al. and VCAM1 from Barron et al. d) Skin-specific nomenclature for fibroblast populations and correspondence to general nomenclature used in main text.

**Extended Data Fig. 3.**
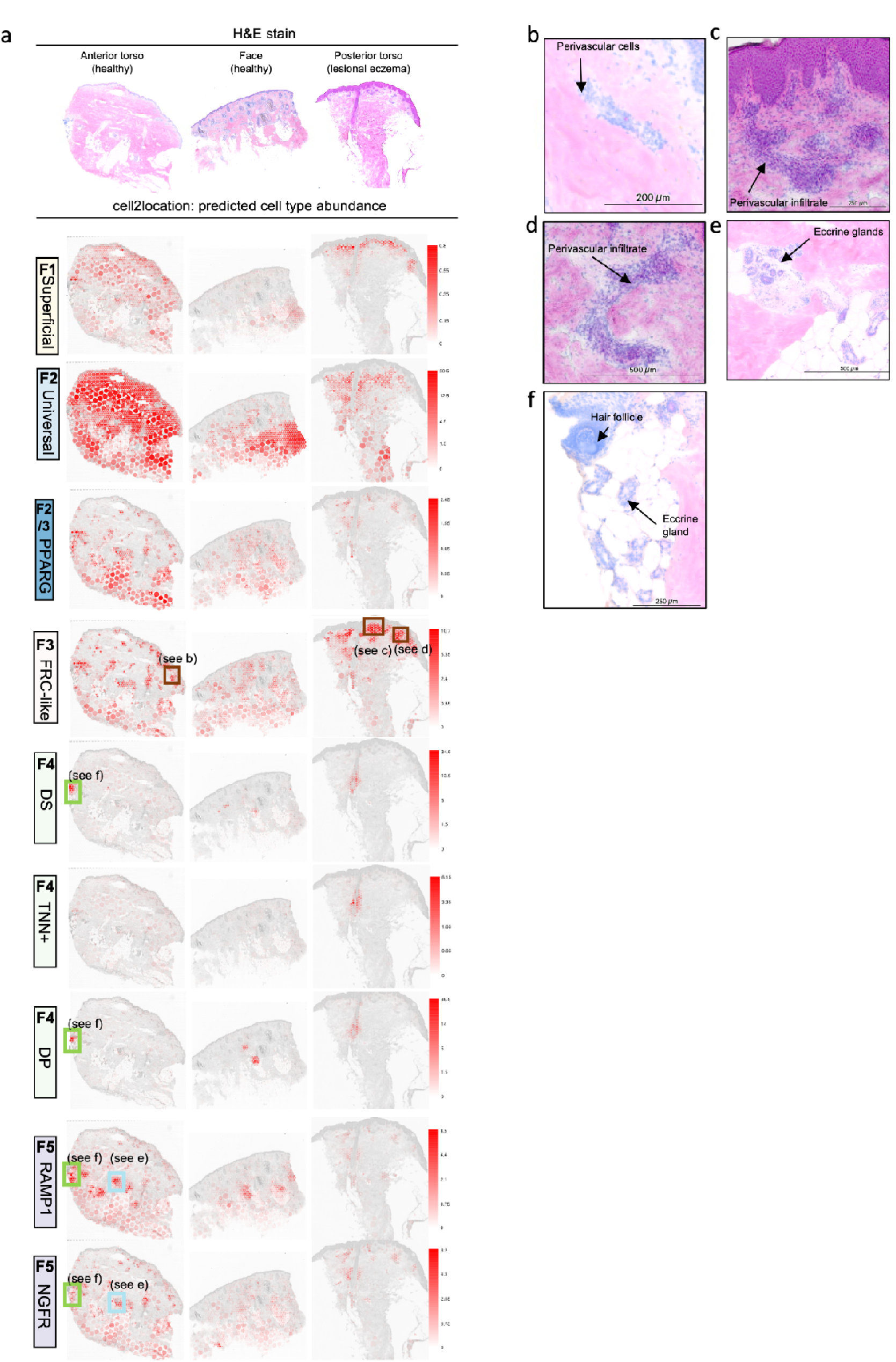
Fibroblast location by deconvolution of fibroblast spots. a) H&E stain of skin slides used for 10x Genomics Visium with cell type abundances predicted by cell2location. For histopathological annotation, each spot represents 55micrometre. H&E: Haematoxylin and eosin b-f) Close-up regions of H&E stained image for regions annotated in panel a.

**Extended Data Fig. 4.**
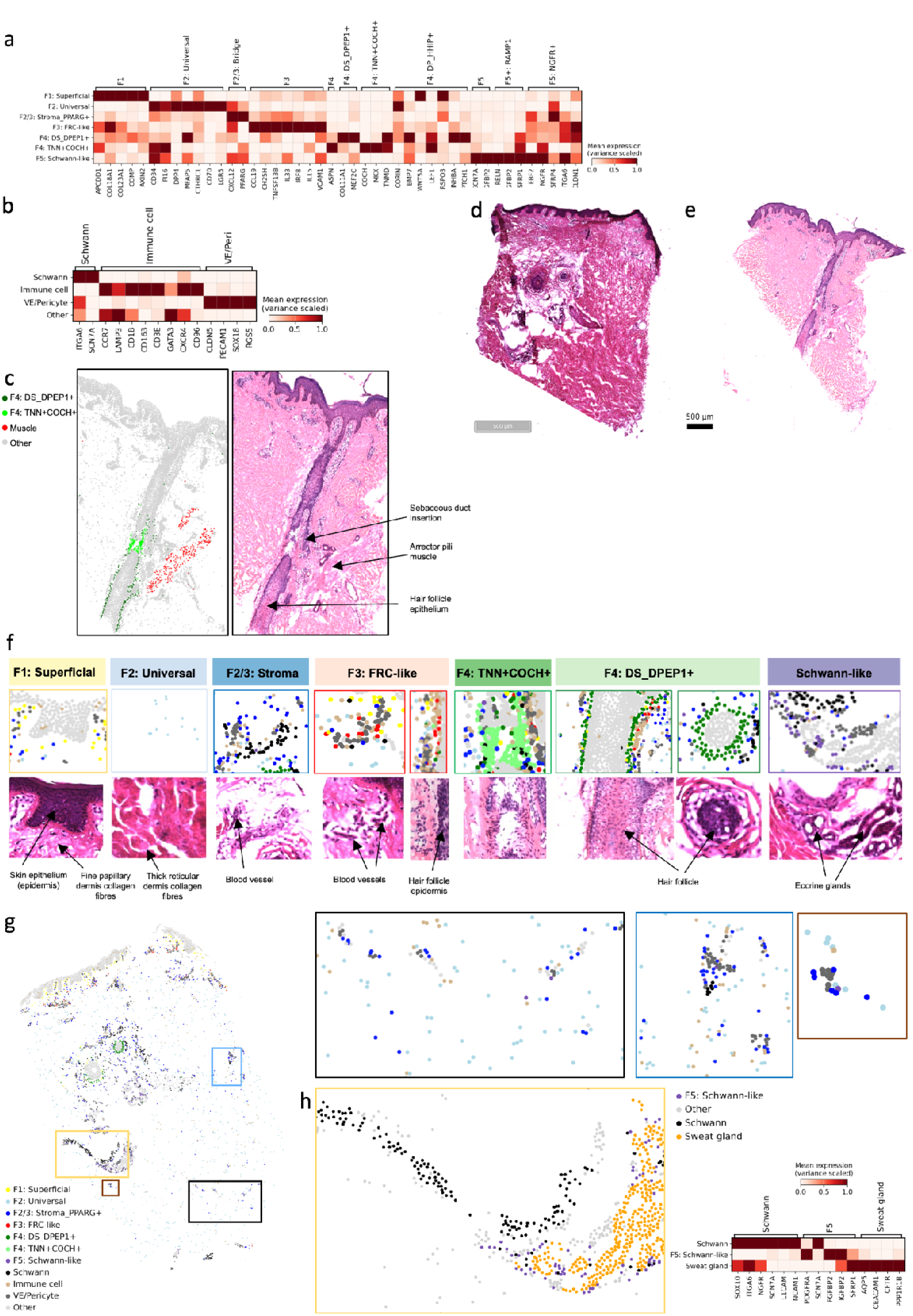
Additional Xenium data. a) Heatmap of marker gene expression in 10x Genomics Xenium data for fibroblast and b) non-fibroblast populations. c) Xenium data coloured by cell type annotation and H&E stain of skin indicating position of *F4: TNN+COCH+,* superior to the insertion of the arrector pili muscle and inferior to the insertion of the sebaceous gland duct. d-e) H&E stain of skin slides used for 10x Genomics Xenium. H&E: Haematoxylin and eosin. f) Summary of fibroblast niches from Fig. 2d but with H&E images adjacent. g) 10x Genomics Xenium of non-lesional atopic dermatitis skin highlighting *F2/3: Stroma_PPARG+* regions and h) *F5: Schwann-like* fibroblasts, including labelled sweat gland cells and heatmap of associated gene expression.

**Extended Data Fig. 5.**
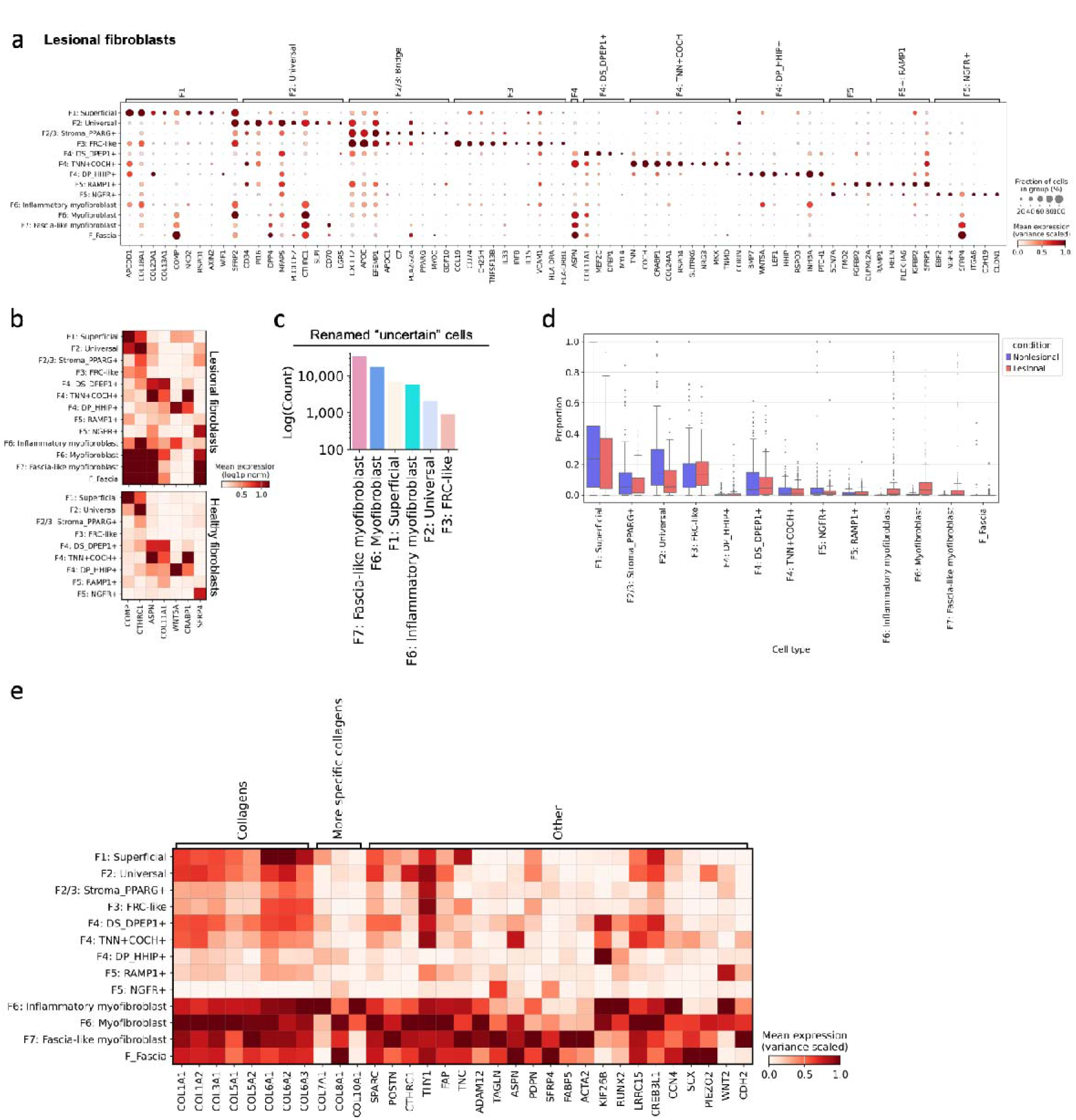
Disease-adapted and disease-specific fibroblast additional data. a) Dotplot of marker genes used for healthy fibroblasts in fibroblasts from lesional/diseased skin. b) Heatmap of marker genes used for healthy fibroblasts that were less specific when diseased fibroblasts were included. c) Top 6 fibroblast populations that were identified from “uncertain” fibroblasts. d) Box plot of fibroblast proportions by site status calculated with scCODA. *F2: Universal* showed depletion and thus was not included as a disease-adapted population. e) Heatmap of gene expression for select collagens, common fibroblast activated genes, and markers we identify for myofibroblasts in skin.

**Extended Data Fig. 6.**
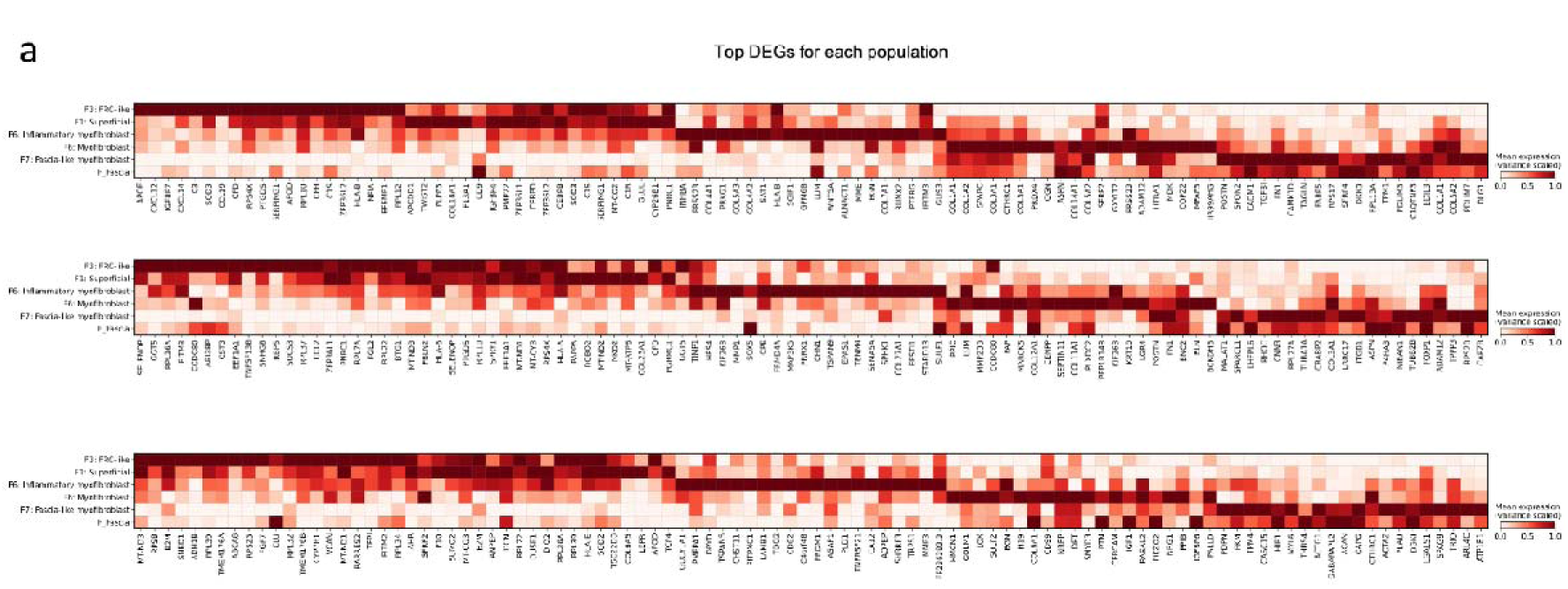
a) Heatmap of top differentially expressed genes (DEGs) for disease-specific and disease-adapted populations. Top row: Top 20 DEGs. Middle row: 20-40 top DEGs. Bottom row: 40-60 top DEGs.

**Extended Data Figure 7.**
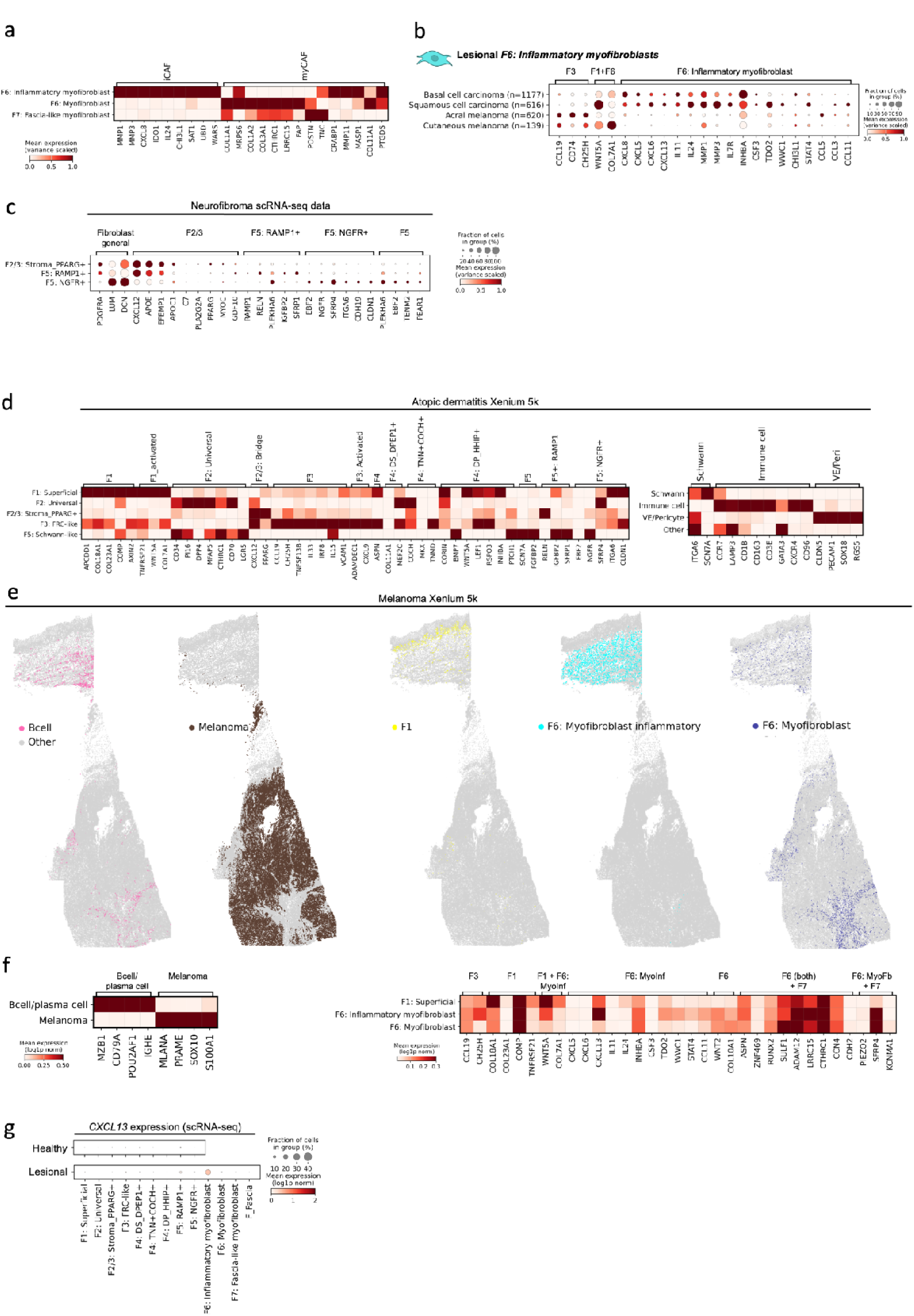
Disease stroma profiles. a) Heatmap of expression of skin cancer-associated fibroblast genes from Forsthuber et al. b) Dotplot of marker genes for inflammatory myofibroblast markers in lesional *F6: Inflammatory myofibroblasts* across different skin cancers. c) Dotplot of marker genes for fibroblasts in neurofibroma. d) Heatmap of marker genes for fibroblast and non-fibroblast populations in Xenium 5k data for lesional/inflamed atopic dermatitis skin. e) Xenium 5k image for cutaneous melanoma (with cells represented by x,y coordinates and coloured by cell type) for single cell types for select populations. f) Heatmap of marker genes for non-fibroblast and fibroblast populations in Xenium 5k data for cutaneous melanoma. g) Dotplot of *CXCL13* expression in fibroblast subtypes from healthy and diseased/lesional skin from scRNA-seq data.

**Extended Data Fig 8.**
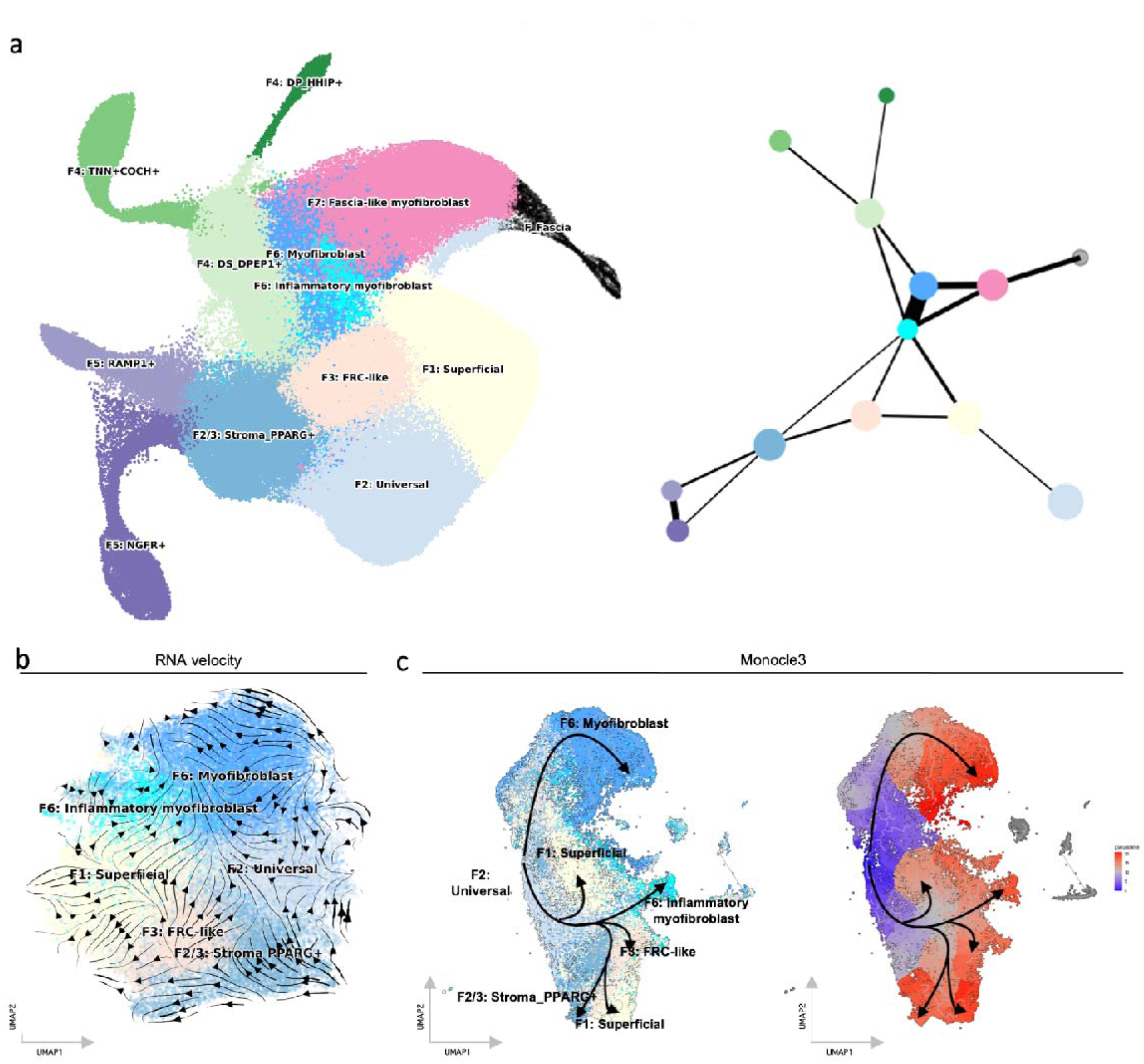
Additional trajectory inference. a) UMAP and partition-based graph abstraction (PAGA) for all lesional and non-lesional fibroblasts. b) RNA velocity (velocity embedding stream) for lesional fibroblasts with velocity data. c) Trajectory inference from Monocle 3 (using *F2: Universal* as the root state) and pseudotime.

**Extended Data Fig 9.**
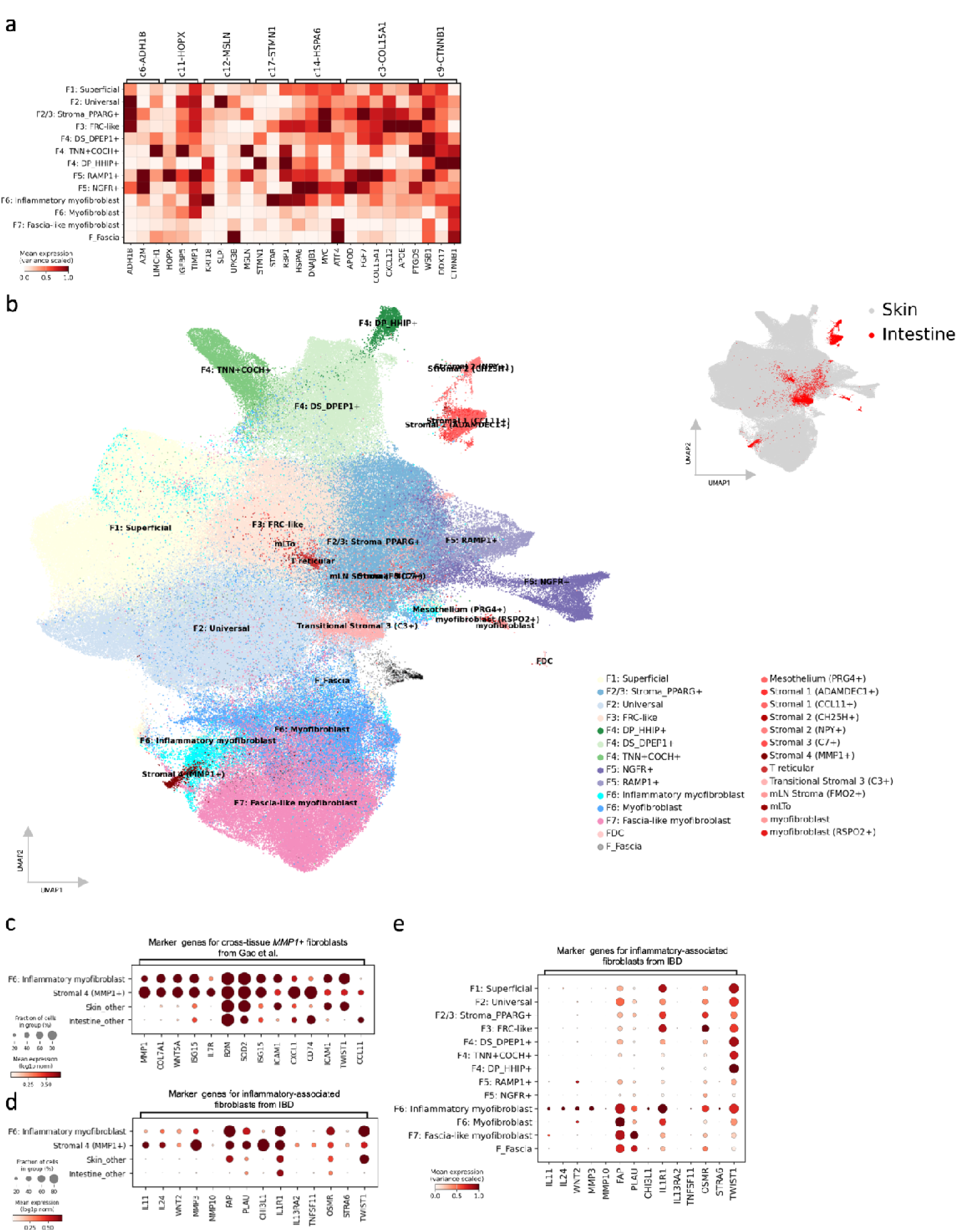
Additional cross-tissue populations and human skin-intestine fibroblast integration. a) Heatmap of marker genes for additional reported shared/universal fibroblast populations from Gao *et al.* b) UMAP visualisation of scVI embedding for skin and intestine fibroblast populations coloured by (left) cell type label, where intestinal populations are in shades of red and (top right) d tissue of origin. c) Dotplot of expression of marker genes for *MMP1+* cross-tissue fibroblasts reported by *Gao et al*. d) Dotplot of expression of marker genes for *inflammatory-associated fibroblasts* reported by *Smillie et al*. IBD: Inflammatory bowel disease. e) Dotplot of marker genes for *inflammatory-associated fibroblasts* reported by *Smillie et al*. in skin fibroblasts, across all skin fibroblast subtypes.

**Extended Data Fig. 10.**
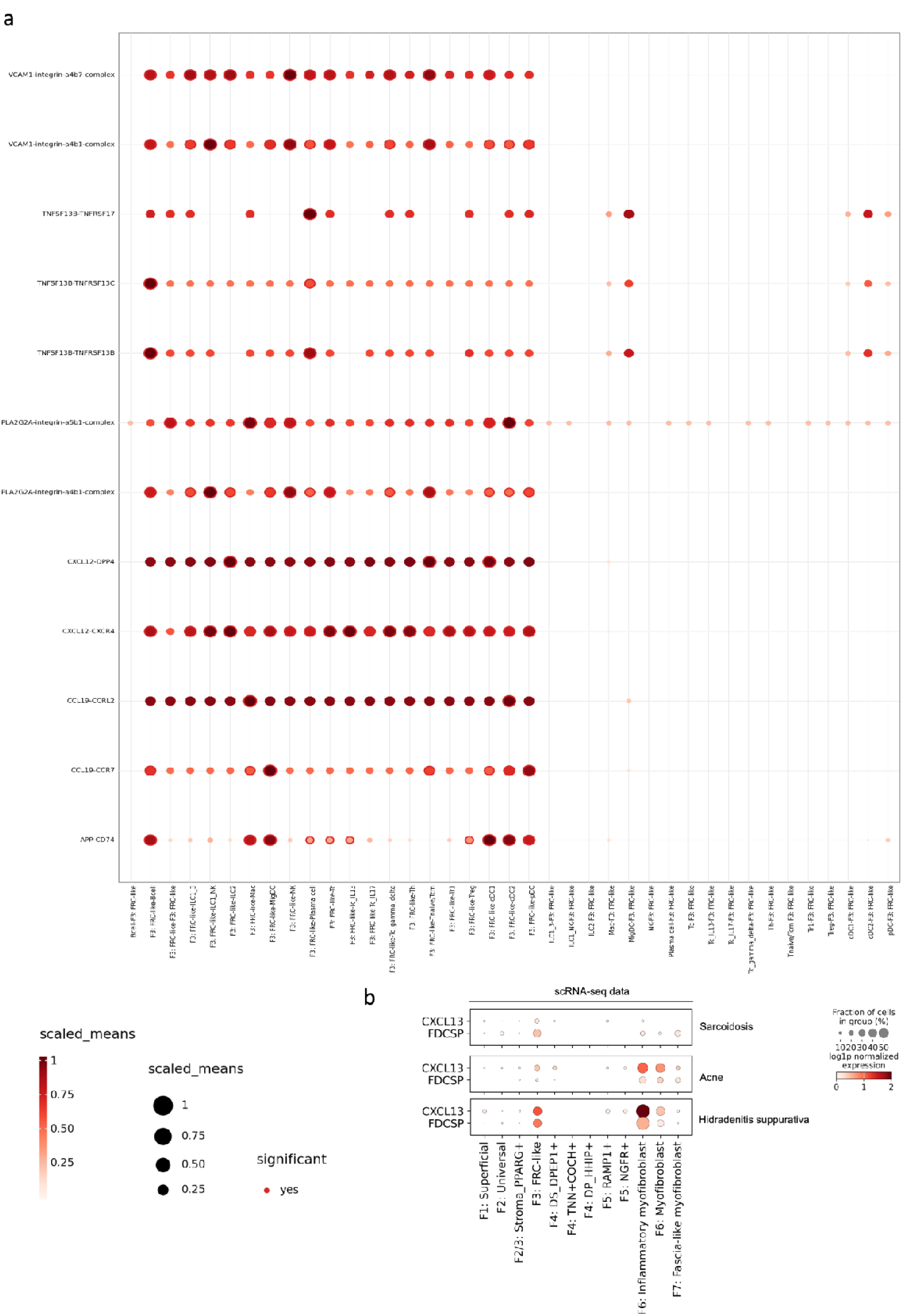
Cell-cell communication results for F3: FRC-like fibroblasts and CXCL13 expression across disease. a) Cell-cell communication results from cellphone DB, including only significant interactions and genes identified as *F3: FRC-like* marker genes previously. b) Dotplot of *CXCL13* expression in hidradenitis suppurativa, acne, and sarcoidosis in lesional fibroblasts.

## Online methods

### Single cell analysis: Atlas integration

Raw scRNA-seq data were downloaded and aligned using STARsolo (GRCh38-2020-A reference). We included publicly-available data generated from fresh skin biopsies using the 10x Genomics Chromium platform. We collected essential metadata (sample id, dataset id, site status, patient status, sex, anatomic location) as more extensive metadata is being collected as part of the Human Skin Atlas. CellBender v.0.3 was used to correct for ambient mRNA^110^. To remove low-quality cells, we included only cells with >200 genes, >1000 and <300 000 total unique molecular identifiers (UMIs), and a mitochondrial gene percentage of <15%. We calculated doublet scores using scrublet on a per sample basis using scrublet and removed cells with a doublet score of >0.35.^111^ Total transcript counts across fibroblast subtypes were similar.

Integration of all cells was performed using single-cell Variational Inference (scVI) using raw counts.^41^ We selected 6000 highly-variable genes (HVGs) as features for the integration. Batch-aware HVG selection was performed by setting the batch key to sample ID. For feature (HVG) selection, we did not include mitochondrial genes; cell cycle genes, from https://github.com/haniffalab/FCA_Whole_Embryo/blob/main/resources/pan_fetal_cc_genes. csv); hypoxic genes; and ribosomal genes. The following hyperparameters were used for the scVI model: number of layers: 2; number of latent dimensions: 30; gene likelihood: zero-inflated negative binomial distribution, and dispersion: gene-batch. An early stopping patience of 5 epochs was used. The batch key was “sampleID” and no other covariates were passed to the model for correction.

Normalisation is a two-step procedure involving depth normalisation and variance stabilisation. We used the shifted logarithm with a scaling factor of 10,000 based on strong performance in a recent benchmarking paper^112^. We constructed a k-nearest neighbours graph (k=30) using the scVI embedding and performed community detection (Leiden algorithm) with resolution 0.1. We selected the fibroblast cluster for further analysis based on canonical marker gene expression (e.g. *PDGFRA*). For a sensitivity analysis, we also selected Schwann cells and pericyte clusters, as well as fascial samples.

### Single cell analysis: Fibroblast-only integration (healthy/non-lesional)

We used the raw counts for healthy and non-lesional fibroblasts as inputs to scVI, using the same parameters as above. We then performed normalisation, kNN construction using the 30-dimension scVI embedding (k=30), and community detection with the Leiden algorithm (resolution=0.3). We also repeated community detection with a higher resolution (1.0) to assess for small, rare but distinct fibroblast subtypes. We manually annotated cell clusters using marker genes selected from prior literature and calculation of differentially expressed genes (DEGs; Extended Data Fig. 2a), which were identified using the t-test with the scanpy rank_genes_groups function. For visualisation in 2 dimensions, we used UMAP with the initialised positions from PAGA implemented in scanpy^113,114^.

### Single cell analysis: Fibroblast-only integration (healthy/non-lesional + disease)

We integrated lesional data with the healthy/non-lesional reference, including fascial fibroblasts, using scPoli^66^. Due to the proposed role of hypoxia in myofibroblast differentiation, we did not exclude hypoxic genes for the lesional integration. We used the following hyperparameters: number of layers: 1; number of latent dimensions: 10; eta(train): 5; eta(query): 10; max epochs(train): 40; max epochs(train): 40. The batch key was “sampleID”. We manually annotated cells labelled as uncertain by performing Leiden clustering and calculating DEGs as above. Visualisation in 2 dimensions was performed using UMAP. Embedding densities were calculated in scanpy using scanpy.pl.embedding_density.

#### Single cell analysis: PROGENy pathway analysis

We used PROGENy pathway analysis to further understand skin fibroblast subtypes in health and disease via the decoupler package^115^. We used the top 500 human genes per pathway and default settings for the multivariate linear model (decoupler.run_mlm).

#### Single cell analysis: scCODA

We used scCODA to plot the distributions (box plots) of fibroblast populations in health and disease^116^.

#### Single cell analysis: Clinical classification

We broadly grouped individual diseases into disease categories (inflammatory + low scarring risk, inflammatory + high scarring risk, cancer, established scarring/fibrosis) based on known disease features. Of note, clear separation of diseases is not possible because of a well-recognised link between inflammation and fibrosis^74^. For example, an inflammatory component to systemic sclerosis is well-recognised and first-line treatments typically include immunosuppressants. Additionally, we included sarcoidosis and granuloma annulare (both disorders of granulomatous inflammation) in ‘inflammatory high-scarring risk’. Most cases of cutaneous sarcoidosis and granuloma annulare do not scar, but fibrosis can occur within granulomas^117^. Additionally, pulmonary sarcoidosis is a well-recognized cause of pulmonary fibrosis. We considered cancer with inflammatory diseases with risk of scarring as fibrosis can occur within lesions (desmoplasia)^118^, and self-resolving melanoma can result in scarring.^119^

For disease proportions, the standard error of the mean (SEM) for each disease category was calculated using the mean proportion for each donor. The SEM for each disease category was derived from the standard deviation of donor-level proportions divided by the square root of the number of donors in that category. This approach quantifies the precision of the mean proportion estimate and accounts for variability across donors.

#### Single-cell: Trajectory inference

For trajectory inference, we used data for which we could calculate RNA velocity using velocyto and scvelo (67 201 lesional fibroblasts)^120,121^. We first used PAGA (as implemented in scanpy),^113^ plotted using a threshold of threshold=0.3, applied to the whole dataset. In future analyses we excluded *F4: Hair-follicle associated* and *F5: Schwann-like fibroblasts,* which appeared to be distinct (Extended Data Fig. 8a), and *F7* fibroblasts and *F_Fascia*, which were observed in few diseases (Fig. 4a).

Velocity pseudotime was calculated using scvelo.^120^ The RNA velocity kernel was calculated using CellRank2. Of note, RNA velocity assumptions such as constant transcription and splicing/degradation rates may be too simple to explain dynamics that arise in multi-lineage cell differentiation^122,123^, and for this reason we use multiple methods (including non-velocity), emphasise our results are predictions, highlight the need for future work, and relate our results to recent findings in animal models.

For Monocle 3 (v 1.3.7)^124^, the expression count matrix along with the corresponding cell and gene metadata from the processed anndata object in scanpy was used to create monocle object (cell_data_set object). The cell_data_set object was then pre-processed using default settings and aligned to correct for batch effects based on the ‘dataset_id’. Dimensionality reduction was performed using ‘UMAP’ as reduction method. Cells were clustered with a resolution of 1[×[10−6 and ’UMAP’ as reduction_method. A trajectory graph was learned by adjusting parameters such as geodesic distance ratio (0.5) and minimal branch length (10) to optimise for large datasets. Finally, cells were arranged in pseudotime by manually selecting root nodes from the *F2: Universal* population. We used *F2: Universal* as the root state in Monocle 3 based on velocity pseudotime results and prior work^4,56^. The ordered and learned graph object was then used to plot the pseudotime trajectory plots.

For inference from methods, we selected results with consistency across different methods. For example, all methods suggested a trajectory from *F2: Universal* to *F6: Myofibroblasts*, and this was a predicted trajectory. However, whereas RNA velocity suggested a trajectory from *F2/3: Stroma_PPARG+* to *F2: Universal*, we did not include this trajectory as directed PAGA revealed an inconsistent result. Through this approach, we aimed to increase the positive predictive value of our results, while acknowledging that this increases the possibility of false negatives.

#### Single-cell: Cross-tissue fibroblast state comparisons

For cross-tissue marker gene comparisons, we selected reported populations from Korsunsky et al., Buecher et al., and Gao et al.^4–6^ As Gao et al. reported 6 universal and 5 shared populations, we showed matching populations in the main figure and other populations in the extended figure. We did not include universal fibroblast populations from these studies (*PI16*, *CD34*, *MFAP5*) as these fibroblasts are well established across tissues^4,5^.

For cross-tissue integrations (adult skin and adult intestine), we concatenated raw count adata objects from intestine^91^ with our data and ran scVI using the same parameters as for the healthy/non-lesonal integration.

#### Single-cell: Prenatal skin-intestine comparison

For prenatal skin and prenatal intestine, we concatenated raw count adata objects from intestine^91^ with our prenatal skin data from a prior publication.^71^ We ran scVI using the same parameters as for the healthy/non-lesonal integration.

#### Single-cell: Mouse FRC-like/Ccl19 comparison

For mouse comparisons, we downloaded the mouse steady state atlas from https://www.fibroxplorer.com/download. We loaded the data as an adata object using pandas2ri in rpy2. We used labels for *Ccl19*+ fibroblast from the original study.

#### Single-cell: cell-cell interactions

We used CellPhoneDB v5 (method 2).^125^ We used our previously published scRNA-seq data from Reynold*s et al*.^31^, restricted to *F3: FRC-like* fibroblasts and immune cells in skin.^31^ We restricted interactions to marker genes for *F3: FRC-like* fibroblasts. We visualized results using ktplotspy.

### Spatial transcriptomics: Visium

We generated new spot-based spatial transcriptomic data using the 10x Genomics Visium Spatial Gene Expression platform from frozen OCT-embedded human adult skin tissue (both healthy and diseased). All research ethics committee and regulatory approval were in place for the collection of research samples at Newcastle and for their storage at the Newcastle Dermatology Biobank (REC reference number: 19/NE/0004). Skin samples were sectioned at 15 µm thickness and the optimal tissue permeabilisation time was determined to be 14 minutes. H&E images were taken using a Zeiss AxioImager with apotome microscope (Carl Zeiss Microscopy) and Brightfield imaging (Zeiss Axiocam 105 48 colour camera module) at 20X magnification. The ZEN blue edition V.3.1 (Carl Zeiss Microscopy) software was used to acquire the H&E images following z-plane and light balance adjustment and image tile stitching. Spatial gene expression libraries were sequenced using an Illumina NovaSeq 6000 to achieve a minimum number of 50,000 read pairs per tissue covered spot.

10x Genomics Visium data was mapped using Spaceranger v.1.3.0 using GRCh38-2020-A reference. Visium provides whole transcriptome coverage over a 55μm diameter spot area. We therefore used the cell2location (v0.1.3) to deconvolute the cell types predicted to be present in a given spot^42^. We constructed a reference signature using sample as batch_key. We included our fibroblast atlas and other skin cell types from Reynolds et al.^31^. Then we performed deconvolution with the following parameters: detection_alpha=20,N_cells_per_location=30, andmax_epochs 50_000. Values of all other parameters were kept default. Following the cell2location tutorial, we used 5% quantile of posterior distribution (q05_cell_abundance_w_sf) as predicted celltype abundances.

### Spatial transcriptomics: Xenium

Non-lesional and lesional eczema human adult skin tissue was used to generate in situ gene expression data using the 10x Genomics Xenium in situ 5k-plex platform. All research ethics committees and regulatory approvals were in place for the collection and storage of research samples at St John’s Institute of Dermatology, Guy’s Hospital, London (REC reference number: EC00/128). Fresh frozen OCT-embedded skin samples were sectioned at 10 µm thickness directly onto the 10X Genomics xenium slide kept at -20°C. We then ran the slides through the 10x Genomics Xenium prime in situ gene expression protocol and imaged using the 10X Genomics Xenium Analyzer. This allowed us to probe 5001 genes at subcellular spatial resolution in human skin sections. We include protein staining to facilitate cell segmentation. Following imaging, the same slides were then used to generate H&E staining and imaged at ×20 magnification on a Hamamatsu Nanozoomer s60.

10x Genomics Xenium data was filtered to exclude cells with <10 genes per cell. We used the shifted log transformation for count normalisation. We calculated the first 50 principal components prior to KNN construction (k=30) and Leiden clustering. We used marker genes from scRNA-seq data to label fibroblast populations through manual annotation. Cell coordinates coloured by cell type were visualised using squidpy (sq.pl.spatial_scatter)^126^.

### Spatial transcriptomics: Xenium niche identification

NicheCompass was run after selecting 1024 spatially variable genes. The number of neighbours selected per cell was 8, and otherwise default settings were used^95^.

## Notes

### Competing Interest Statement

In the past 3 years, S.A.T. has consulted or been a member of scientific advisory boards at Roche, Genentech, Biogen, GlaxoSmithKline, Qiagen and ForeSite Labs and is an equity holder of Transition Bio and EnsoCell.

